# CLASP2 promotes repair of kinesin-1 damage to the microtubule lattice

**DOI:** 10.64898/2026.06.29.735199

**Authors:** Jakia Jannat Keya, Rahul Riberio, Yang Yue, Elizabeth J. Lawrence, Marija Zanic, Kristen J. Verhey

**Affiliations:** Department of Cell and Developmental Biology, University of Michigan, Ann Arbor, MI, USA; Department of Biophysics, University of Michigan, Ann Arbor, USA; Department of Cell and Developmental Biology, Vanderbilt University School of Medicine, Nashville, TN, USA

## Abstract

Microtubules are cytoskeletal polymers that play essential roles in eukaryotic cells, including structural support, cell division, and intracellular transport. During intracellular transport, kinesin motor proteins move cargo along microtubule tracks via their processive stepping. Recent studies have shown that the kinesin-1 KIF5C can damage the microtubule lattice while stepping. Microtubule damage can be repaired through incorporation of new tubulin subunits, however, excessive lattice damage results in microtubule breakage and disassembly. To identify cellular factors involved in microtubule repair, we performed an siRNA screen targeting microtubule-associated proteins (MAPs) known to regulate microtubule dynamics and stability. Based on the results, we investigated whether the end binding protein EB1 and cytoplasmic linker-associated protein 2 (CLASP2) contribute to repair of microtubule damage. To test this, we used a microtubule destruction assay in which damage was induced in microtubules gliding over surfaces coated with wild-type or mutant KIF5C proteins. Our findings suggest that CLASP2 directly facilitates microtubule repair, whereas EB1 does not. We further examined CLASP function using a microtubule repair assay and found that CLASP2 promotes repair by enhancing tubulin incorporation and reducing microtubule breakage. Together, these findings demonstrate that CLASP proteins play an important role in repairing and protecting against lattice damage caused by kinesin-1 motor activity. Our results further suggest that MAPs can directly regulate microtubule lattice integrity under mechanical stress generated by motor protein-driven intracellular transport.

## Introduction

Microtubules are a fundamental component of the eukaryotic cytoskeleton, playing key roles in cell division, cell structure and organization, and intracellular transport. They are hollow cylindrical polymers that self-assemble from tubulin subunits (heterodimers of α-tubulin and β-tubulin) and exhibit energy-dependent, dissipative growth dynamics [1–5]. During polymerization, GTP-bound tubulins assemble into a stable microtubule structure. Once incorporated, β-tubulin hydrolyzes GTP to GDP resulting in GDP-bound lattice. A “GTP cap” at the growing plus end stabilizes the microtubule, but if it is lost, the microtubule undergoes rapid depolymerization. A rescue occurs when shrinkage switches back to growth due to addition of new GTP-tubulin subunits or encountering GTP-rich islands in the lattice. In cells, the minus end is often anchored whereas the plus end is highly dynamic and undergoes most growth and shrinkage.

Recent work has demonstrated that tubulin can also undergo nucleotide-dependent exchange along the microtubule lattice. Tubulin dissociation from the lattice can occur due to thermal fluctuations, local mechanical forces, the activity of severing enzymes, and the stepping of motor proteins [6–9]. Kinesins are a superfamily of motor proteins that drive the transport of cargoes during intracellular trafficking [10–12]. Traditionally, microtubules have been viewed simply as tracks for kinesin motility [13,14]. However, recent studies have revealed that kinesin-1 can cause tubulin dissociation from the microtubule lattice, *i*.*e*. damage sites, while stepping [15–17]. Further, a mutant version of the kinesin-1 KIF5C causes increased lattice damage compared with wild-type (WT) kinesin-1 in cells assays [17]. Although soluble tubulin can repair lattice defects induced by kinesin-1 motor stepping [15–17], damage that exceeds the microtubules’ capacity for self-repair results in microtubule breakage and disassembly [17, 18]. These findings suggest that cellular factors contribute to the recognition and repair of microtubule damage.

Several findings suggest that lattice repair may be facilitated by proteins known to regulate microtubule growth and shrinkage at the plus end. The end-binding (EB) proteins are central regulators of microtubule plus end dynamics. They can directly regulate persistent microtubule nucleation and growth, promoting both catastrophe and rescue in vitro [19–22]. That EB proteins could play a role in lattice repair is suggested by their colocalization with newly-incorporated tubulin at damage sites induced by severing enzymes or laser light [23–25]. Cytoplasmic Linker Protein-170 (CLIP-170) is an EB-associated protein that facilitates rescue of microtubule plus end growth in cells and in vitro [26, 27]. CLIP-170 is thought to facilitate rescue of microtubule growth via its two cytoskeleton-associated protein glycine-rich (CAP-Gly) domains that bind soluble tubulin [28–30]. That CLIP-170 could play a role in lattice repair is suggested by the finding that in cells, CLIP-170 localizes along the lattice at GTP-like tubulin islands that are hotspots for rescue events [31, 32]. CLASPs are highly conserved EB-and CLIP-170-interacting proteins that play a critical role in stabilizing microtubules by preventing catastrophes and facilitating rescues [33–38]. CLASPs contain tumor overexpressed gene (TOG) domains which enable binding to both the microtubule lattice and soluble tubulin [34, 38–43]. That CLASPs could play a role in lattice repair is suggested by findings that CLASP2α and its TOG2 domain can stimulate tubulin incorporation at laser-induced damage sites and thereby increase the efficiency of lattice repair [39]. The microtubule-associated protein (MAP) Tau also facilitates repair by increasing tubulin exchange at defect sites, challenging the view of tau as a passive stabilizer [44].

Whether these proteins play a role in the repair of motor-induced damage of the microtubule lattice has not been tested. Here we investigated the effect of various MAPs on the repair of microtubule damage caused by the kinesin-1 KIF5C. We first used siRNA-mediated knockdown of various MAPs to test their ability to facilitate lattice repair in cells overexpressing kinesin-1. We found that knockdown of EB1, DCTN1, or CLASP2 resulted in increased microtubule breakage and fragmentation, suggesting that these proteins may play a role in lattice repair. We then used *in vitro* reconstitution assays to directly test the roles of EB1 and CLASP2γ in lattice repair and microtubule stabilization. We first used a microtubule destruction assay in which KIF5C stepping induces lattice damage that leads to loss of microtubules over time. We found that addition of CLASP2γ but not EB1 was able to suppress the loss of microtubules. We then used a microtubule repair assay to investigate how CLASP2γ prevents microtubule destruction. We found that CLASP2γ promotes the incorporation of free tubulin along the microtubule lattice. These results suggest that CLASP proteins are critical factors for protecting and repairing lattice damage caused by kinesin-1 stepping along the microtubule. Our results suggest that cells can repair microtubule damage through the action of MAPs such as CLASP to protect microtubule from the mechanical stress of kinesin-1 stepping.

## Results

### siRNA screen suggests roles for EB1 and CLASP2 in repair of KIF5-induced microtubule damage

To investigate the contribution of various MAPs to the repair of microtubule damage induced by kinesin-1 stepping, we used siRNA to knockdown the expression of various MAPs. We hypothesized that the loss of a repair factor would result in extensive microtubule damage and fragmentation in cells expressing KIF5C. We transfected hTERT-RPE1 cells with siRNA pools targeting various MAPs or with no siRNA or scramble siRNA controls. After 24 hr, the cells were transfected with plasmids for expression of constitutively-active KIF5C(1-560) [hereafter KIF5C(WT)] or the mutant version KIF5C(1-560,Δ6) [hereafter KIF5C(Δ6)] which causes increased damage to the lattice and extensive microtubule breakage and fragmentation in cells [17]. After an additional 24 hr, the cells were fixed and stained to visualize the microtubule network. We applied a qualitative grading of the microtubule network in each cell (Fig. 1B), scoring each cell as having normal microtubules (straight microtubules with no extensive curling or breakage), medium damage (microtubules have some curling and/or fragmentation, particularly at the cell periphery), or high damage (microtubules display extensive fragmentation and retraction from the cell edge) (Fig. 1A, 1B).

**Figure 1.**
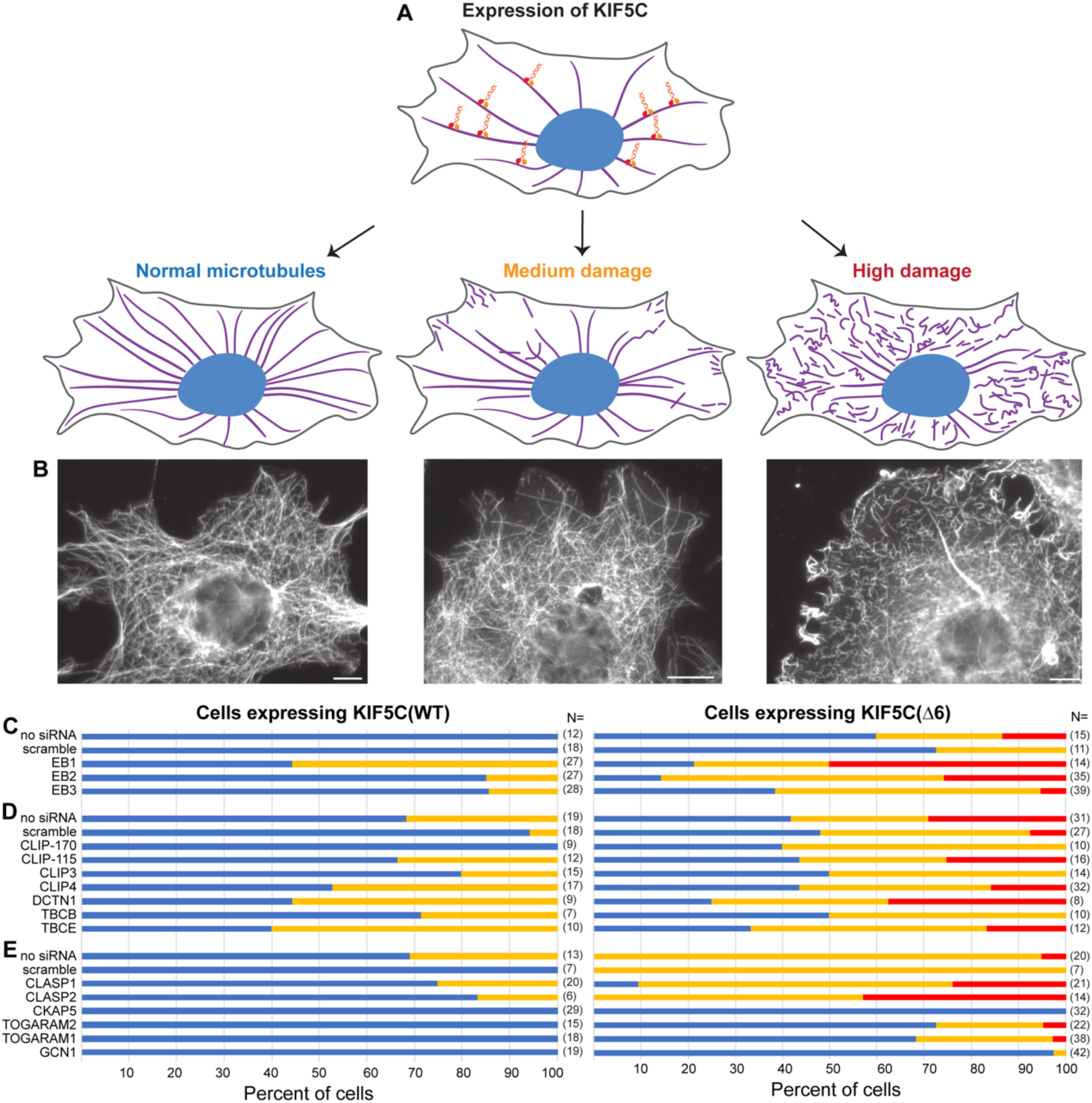
siRNA screen for MAPs that influence kinesin-induced microtubule damage and destruction in cells. (A) Schematic of experiment. hTERT-RPE1 cells were transfected with siRNA and then with plasmids for expression of KIF5C protein. After 24 h, the cells were fixed and stained with an antibody against tubulin and the resulting microtubule breakage and fragmentation per cell was qualitatively scored as normal (no damage), medium damage, or high damage. (B) Representative images of microtubule destruction. Scale bars: 10 µm. (C) Microtubule damage in cells where EB1, EB2 or EB3 was knocked down in cells expressing (left) KIF5C(WT) or (right) KIF5C(Δ6) motors. N = number of cells analyzed across three experiments. (D) Microtubule damage in cells where the indicated CAP-Gly domain-containing proteins were knocked down in cells expressing (left) KIF5C(WT) or (right) KIF5C(Δ6) motors. N = number of cells analyzed across six experiments (no siRNA, scramble, CLIP-170, CLIP-115, CLIP3, CLIP4) or three experiments (DCTN1, TBCB, TBCE). (E) Microtubule damage in cells where the indicated TOG domain-containing proteins were knocked down in cells expressing (left) KIF5C(WT) or (right) KIF5C(Δ6) motors. N = number of cells analyzed across three experiments (no siRNA, scramble, CLASP1, CLASP2) or four experiments (CKAP5, TOGARAM2, TOGARAM1, GCN1).

We first tested whether EB proteins play a role in the repair of KIF5-induced damage to the microtubule lattice. Knockdown of EB1 had the greatest effect and resulted in an increased number of cells with a medium level of microtubule damage in cells expressing KIF5C(WT) compared to the no siRNA and scramble siRNA control conditions (Fig. 1C, left). Furthermore, knockdown of EB1 resulted in an increase in cells with high levels of microtubule damage in cells expressing KIF5C(Δ6) compared to the control conditions (Fig. 1C, right).

We then tested whether proteins containing a tubulin-binding CAP-Gly domain, such as CLIP-170, are involved in repair of KIF5C-induced damage. Surprisingly, knockdown of CLIP-170 resulted in less microtubule damage in cells expressing KIF5C(WT) or KIF5C(Δ6) compared to the control conditions or the other siRNA conditions (Fig. 1D). In contrast, loss of the other CAP-Gly containing proteins resulted in similar or more damage than the control conditions, particularly knock-down of DCTN1, TBCE, or CLIP4 (Fig. 1D).

Next, we tested whether proteins that contain a TOG domain, such as CKAP5 (ch-TOG) or the CLASPs, are involved in repair of KIF5C-induced microtubule damage. Knockdown of CLASP1 or CLASP2 had minimal effect in cells expressing KIF5C(WT) but caused increased microtubule damage in cells expressing KIF5C(Δ6) (Fig. 1E).

Based on these results, we conclude that loss of EB1, DCTN1 or TBCE results in the highest levels of microtubule damage in cells expressing KIF5C(WT) whereas loss of EB1, DCTN1 or CLASP2 results in the highest levels of microtubule damage in cells expressing KIF5C(Δ6). Given these results and the previous reports on localization of EB and CLASP proteins to damage sites [23–25, 39], we chose to directly test the roles of EB1 and CLASP2 in repair of KIF5C-induced microtubule damage.

### EB1 does not play a direct role in repairing KIF5-induced microtubule damage

To directly test the roles of EB1 and CLASP2 in lattice repair, we used a microtubule destruction assay, a modified microtubule gliding assay in which KIF5C stepping results in lattice damage such that gliding microtubules break and depolymerize over time. We purified KIF5C(WT)-Halo-TwinStrep and KIF5C(Δ6)-Halo-TwinStrep proteins from Sf9 cells (Fig. S1). Microtubules were polymerized in the presence of GTP to generate GDP-microtubules and stabilized by 25% glycerol. Microtubule destruction assays were prepared in which KIF5C-Halo-TwinStrep motors were attached to streptavidin-coated flow cells and then glycerol-stabilized GDP-microtubules were introduced (Fig. S2A). Upon addition of ATP, motor stepping causes damage to the lattice while gliding the microtubules. The amount of damage and destruction can be tuned by the amount of free tubulin. At 7 µM free tubulin (unlabeled), microtubule destruction occurred more rapidly for KIF5C(τι6) than KIF5C(WT) (Fig. S2B, S2C, Movie 1, Movie 2), consistent with previous work [17].

We thus used the microtubule destruction assay to investigate the effect of EB1 on repair of KIF5C-induced lattice damage. EB1-EGFP was purified from bacterial cells (Fig. S1) and added to microtubule destruction assays at 50 nM or 100 nM based on previous studies [23, 24, 45]. Flow chambers were incubated sequentially with BSA-biotin, streptavidin, and KIF5C(WT) or KIF5C(Δ6) motors. Glycerol-stabilized GDP-microtubules were then added in the presence of 1 mM GTP, 2 mM ATP, 7 µM free tubulin, and different concentrations of EB1-EGFP (Fig. 2A, 2B). We found that the addition of EB1-EGFP did not block the destruction of microtubules, rather, the presence of EB1-GFP resulted in a small but statistically significant decrease in the number of microtubules remaining after damage induced by KIF5C(WT) and KIF5C(Δ6) (Fig. 2C, Fig. S3). We conclude that EB1 does not have a direct role in repairing lattice damage caused by the stepping of kinesin-1.

**Figure 2.**
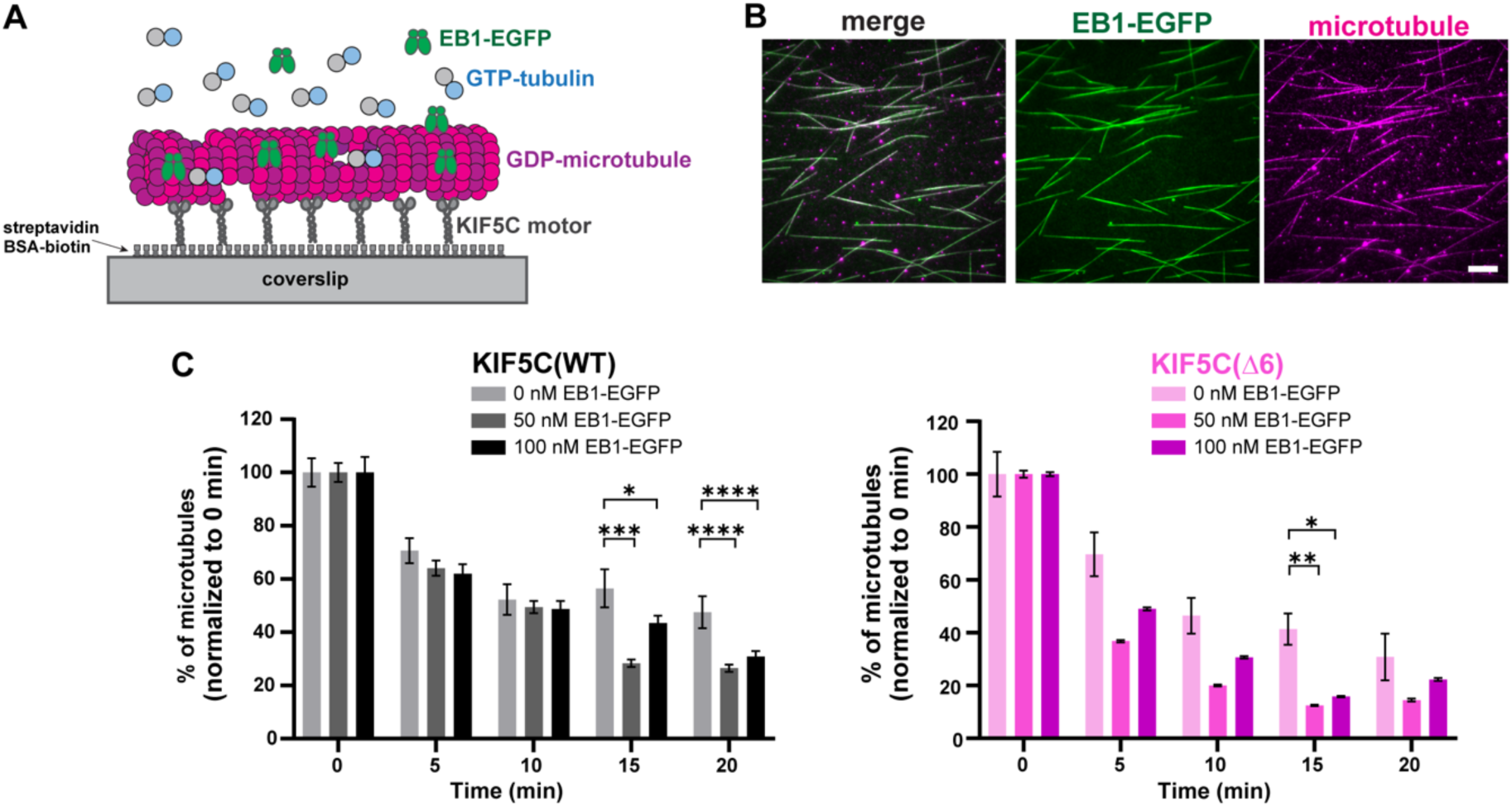
EB1 does not protect microtubules from kinesin-induced destruction. (A) Schematic of microtubule destruction assay. In this modified microtubule gliding assay, glycerol-stabilized GDP-microtubules are driven by KIF5C motors in the presence of 2 mM ATP, 1 mM GTP and 7 µM unlabeled free tubulin and in the absence or presence of EB1-EGFP. (B) Representative image at 0 min time point of assay with 100 nM EB1-GFP (green) and GDP-microtubules (magenta). Scale bar: 10 µm. (C) Quantification of microtubule destruction over time for (left) KIF5C(WT) or (right) KIF5C(Δ6) in the absence or presence of the indicated concentrations of EB1-EGFP. The total number of microtubules per field of view (FOV) was quantified at 0 min (immediately after ATP addition to start KIF5C motility), 5, 10, 15, and 20 min. The percent of microtubules remaining in each FOV was determined, averaged for 27 FOV across three independent experiments, and normalized to the 0 min time point (error bars: SE). * p<0.05, ** p<0.01, *** p<0.001, **** p<0.0001 (one-way Anova test).

### CLASP2γ prevents microtubule damage induced by KIF5C motors in microtubule destruction assay

We then investigated the role of CLASP2γ-EGFP in the repair of KIF5C-induced lattice damage. CLASP2γ-EGFP-StrepII was purified from Sf9 cells (Fig. S1) and added to microtubule destruction assays at 100 or 200 nM based on previous studies [34, 46]. As the StrepII tag would bind to streptavidin-coated flow cells (Fig. 2A), we switched to neutravidin-coated flow cells (Fig. 3A) for these experiments as StrepII shows little binding to neutravidin surfaces [47, 48]. Then, to attach KIF5C motors to the neutravidin-coated surfaces, we used Avi-tagged KIF5C biotinylated by co-expression with BirA in COS-7 cells and added to the reaction in cell lysates as we have done previously [17]. Thus, flow chambers were incubated sequentially with BSA-biotin, neutravidin, and then KIF5C-AviTag motors. Glycerol-stabilized GDP-microtubules were then added in the presence of 1 mM GTP, 2 mM ATP, 7 µM free tubulin, and various concentrations of CLASP2γ-EGFP (Fig. 3A, 3B).

**Figure 3.**
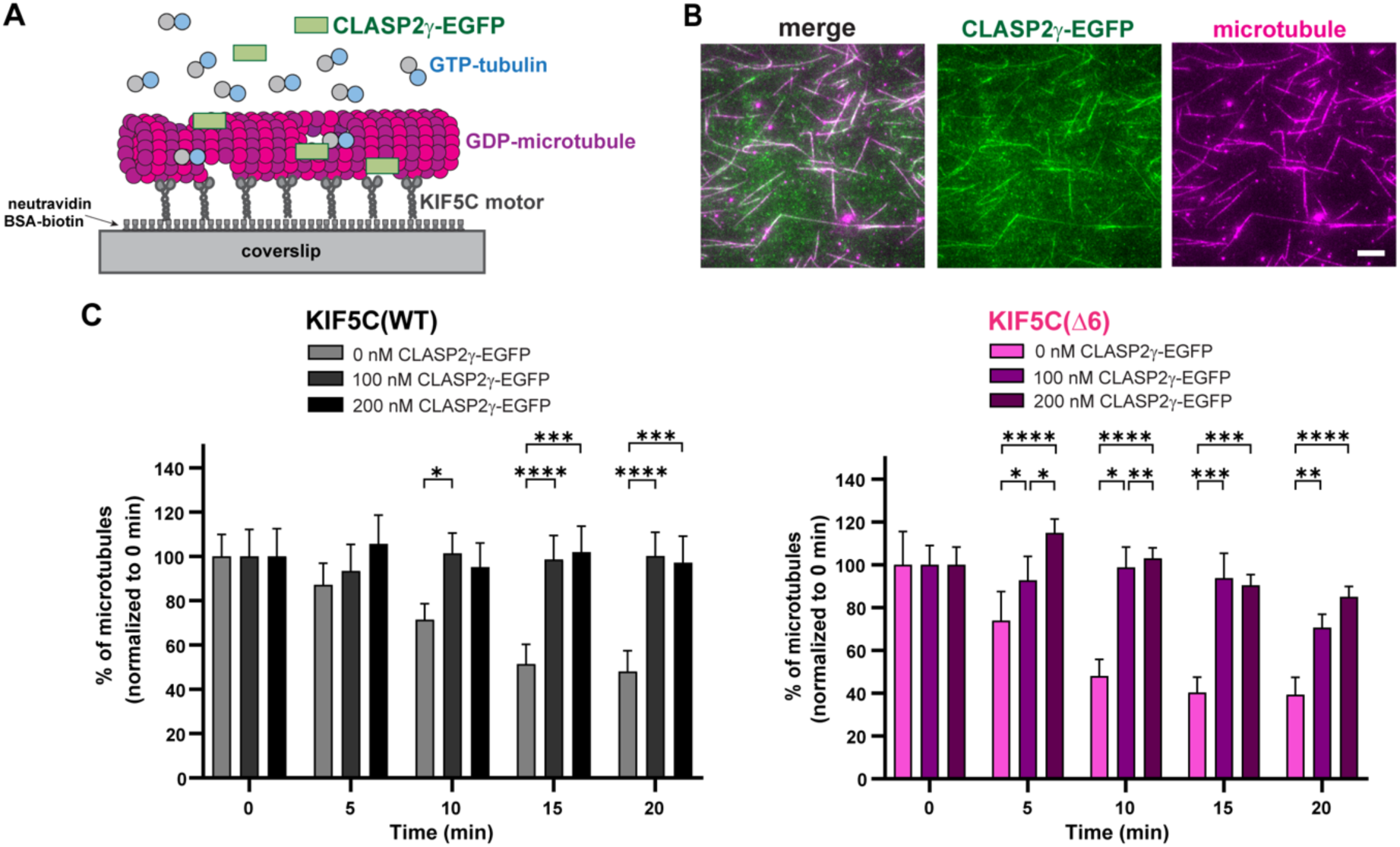
CLASP2γ protects microtubules from kinesin-induced destruction. (A) Schematic of microtubule destruction assay. Glycerol-stabilized GDP-microtubules are driven by KIF5C motors in the presence of 2 mM ATP, 1 mM GTP and 7 µM unlabeled free tubulin in the absence or presence of CLASP2γ-EGFP. (B) Representative images at 0 min time point of 200 nM CLASP2γ-EGFP (green) and GDP-microtubules (magenta). Scale bar: 10 µm. (C) Quantification of microtubule destruction over time for (left) KIF5C(WT) or (right) KIF5C(Δ6) in the absence or presence of the indicated concentrations of CLASP2γ-EGFP. The total number of microtubules per field of view (FOV) was quantified at 0 min (immediately after ATP addition to start KIF5C motility), 5, 10, 15, and 20 min. The percent of microtubules remaining in each FOV was determined, averaged for 27 FOV across three independent experiments, and normalized to the 0 min time point (error bars: SE). * p<0.05, ** p<0.01, *** p<0.001, **** p<0.0001 (one-way Anova test).

We found that the addition of CLASP2γ-EGFP prevented the destruction of microtubules due to damage induced by both KIF5C(WT) and KIF5C(Δ6) motors (Fig. 3C, Fig. S4). For lattice damage induced by KIF5C(WT), the addition of 100 nM CLASP2γ-EGFP was sufficient to prevent microtubule destruction at all time points. After 20 min of microtubule gliding, 100% of the microtubules remained in the presence of 100 nM CLASP2γ-EGFP whereas only 48% remained in the absence of CLASP2γ-EGFP (Fig. 3C). For KIF5C(Δ6), addition of CLASP2γ-EGFP partially prevented microtubule destruction. After 20 min of microtubule gliding, 70% of the microtubules remained in the presence of 100 nM CLASP2γ-EGFP and 85% remained in the presence of 200 nM CLASP2γ-EGFP whereas only 39% remained in the absence of CLASP2γ-EGFP (Fig. 3C). These results indicate that CLASP2γ plays a direct role in preventing the destruction of microtubules damaged by the stepping of kinesin-1 motors.

### CLASP2γ repairs KIF5C-induced damage by facilitating free tubulin incorporation into the lattice

To understand how CLASP2γ prevents the loss of microtubules in the microtubule destruction assay, we considered the possibility that CLASP2γ simply limits kinesin-1 stepping. While this seemed unlikely given the ability of kinesin-1 to drive microtubule gliding in the presence of CLASP2γ-EGFP (Fig. 3A), we directly measured the effects of CLASP2γ-EGFP on kinesin-1 stepping in a single-molecule motility assay (Fig. 4A). Purified KIF5C(Δ6)-Halo^554^-FLAG motors (Fig. S1) were added to flow chambers containing taxol-stabilized microtubules in the absence or presence of CLASP2γ-EGFP, and their motility properties were determined from kymographs (Fig. 4A, 4B). Little to no change was observed for the landing rate (Fig. 4C) or the velocity (Fig. 4D) of KIF5C(Δ6)-Halo^554^-FLAG motors in the presence of 200 nM CLASP2γ-EGFP, however, the run length decreased significantly in presence of CLASP2γ-EGFP (median 2.0 µm) compared to in the absence of CLASP2γ-EGFP (median 4.8 µm) indicating that CLASP2γ reduces the processivity of KIF5C(Δ6) on microtubules (Fig. 4E). These results suggest that CLASP2γ does not impair KIF5C’s interaction with microtubules but can serve as an obstacle to limit its processivity.

**Figure 4.**
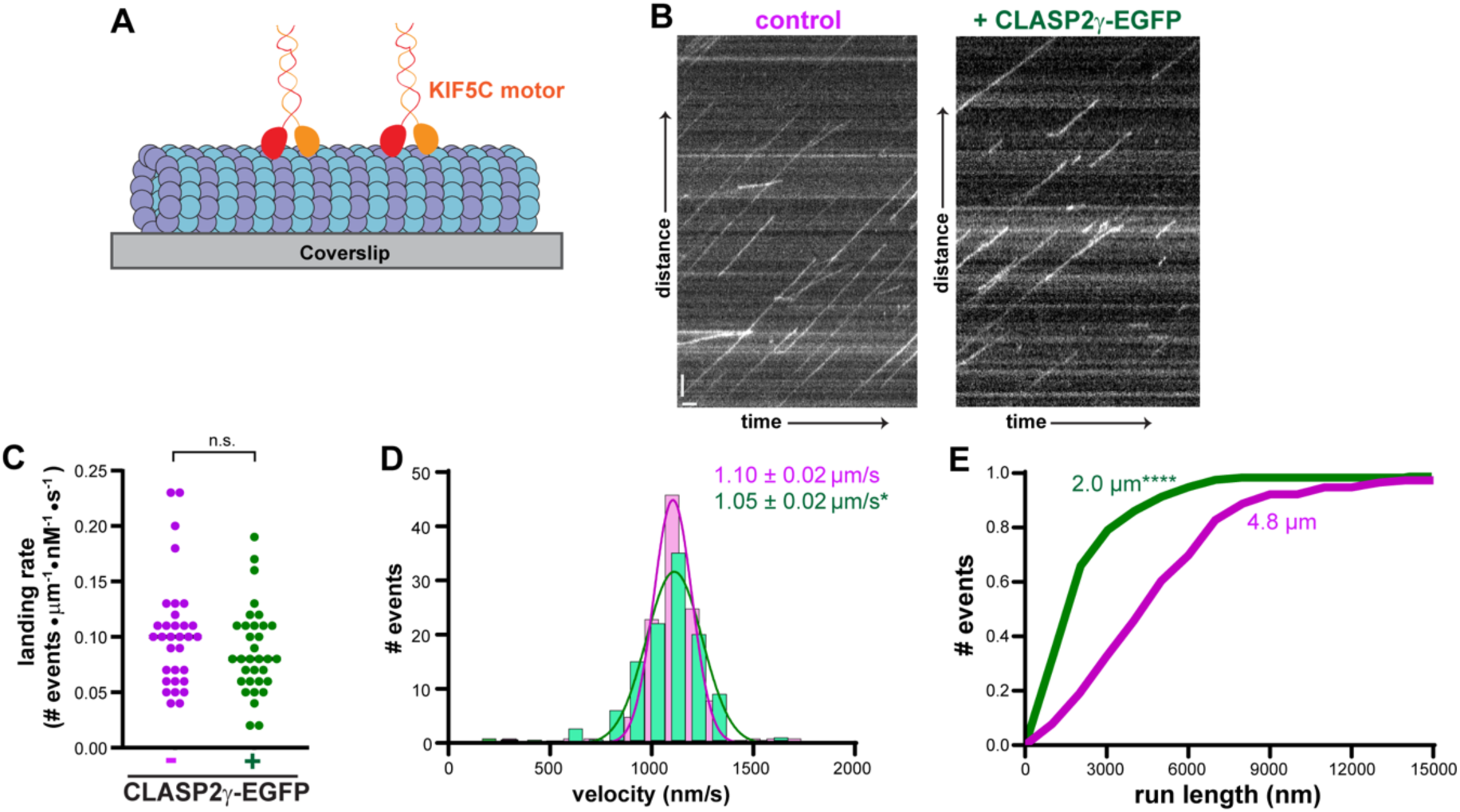
KIF5C(Δ6) shows reduced processivity in the presence of CLASP2γ. (A) Schematic of single-molecule motility assay. The motility of KIF5C(Δ6) along taxol-stabilized microtubules was determined in the absence or presence of 200 nM CLASP2γ-EGFP. (B) Representative kymographs with time displayed on the x axis (scale bar: 2 s) and distance displayed on the y axis (scale bar: 2 µm). (C) KIF5C(Δ6) landing rate in absence or presence of CLASP2γ-EGFP. N = 32 microtubules across three independent experiments; n.s: not significant (two-tailed t test). (D,E) Motility properties in the absence or presence of CLASP2γ-EGFP. N = 115 events for 0 nM and 112 events for 200 nM CLASP2γ-EGFP across three independent experiments. (D) KIF5C(Δ6) velocity plotted as histograms with the mean ± SD indicated at the top right of the graph for KIF5C(Δ6) motility in absence (magenta) or presence of 200 nM CLASP2γ-EGFP (green). *p < 0.05 (two-tailed Welch’s t test). (E) KIF5C(Δ6) run length plotted as cumulative distribution functions with median run length of 4.8 µm in the absence of CLASP2γ-EGFP (magenta) and 2.0 µm in the presence of 200 nM CLASP2γ-eGFP (green). ****p < 0.0001 (Kolmogorov-Smirnov test).

We next considered the possibility that CLASP2γ limits microtubule destruction by preventing the loss of tubulin from the microtubule lattice. To test this, we performed lattice repair assays in which the incorporation of free, fluorescent tubulin into the microtubule lattice is measured after kinesin-1 stepping. We generated GDP-lattice microtubules (grown from GMPCPP-seeds and stabilized with GMPCPP-caps) and added purified KIF5C(Δ6)-Halo-TwinStrep motors and 10 µM soluble Hilyte647-labeled tubulin in the absence or presence of 200 nM CLASP2γ-EGFP-StrepII (Fig. 5A). We found that regions of free tubulin incorporation into the GDP-microtubule lattice (*i*.*e*. repair sties) could be detected both in the absence and presence of CLASP2γ-EGFP (Fig. 5B). This result suggests that CLASP2γ does not simply prevent loss of tubulin from the lattice.

**Figure 5.**
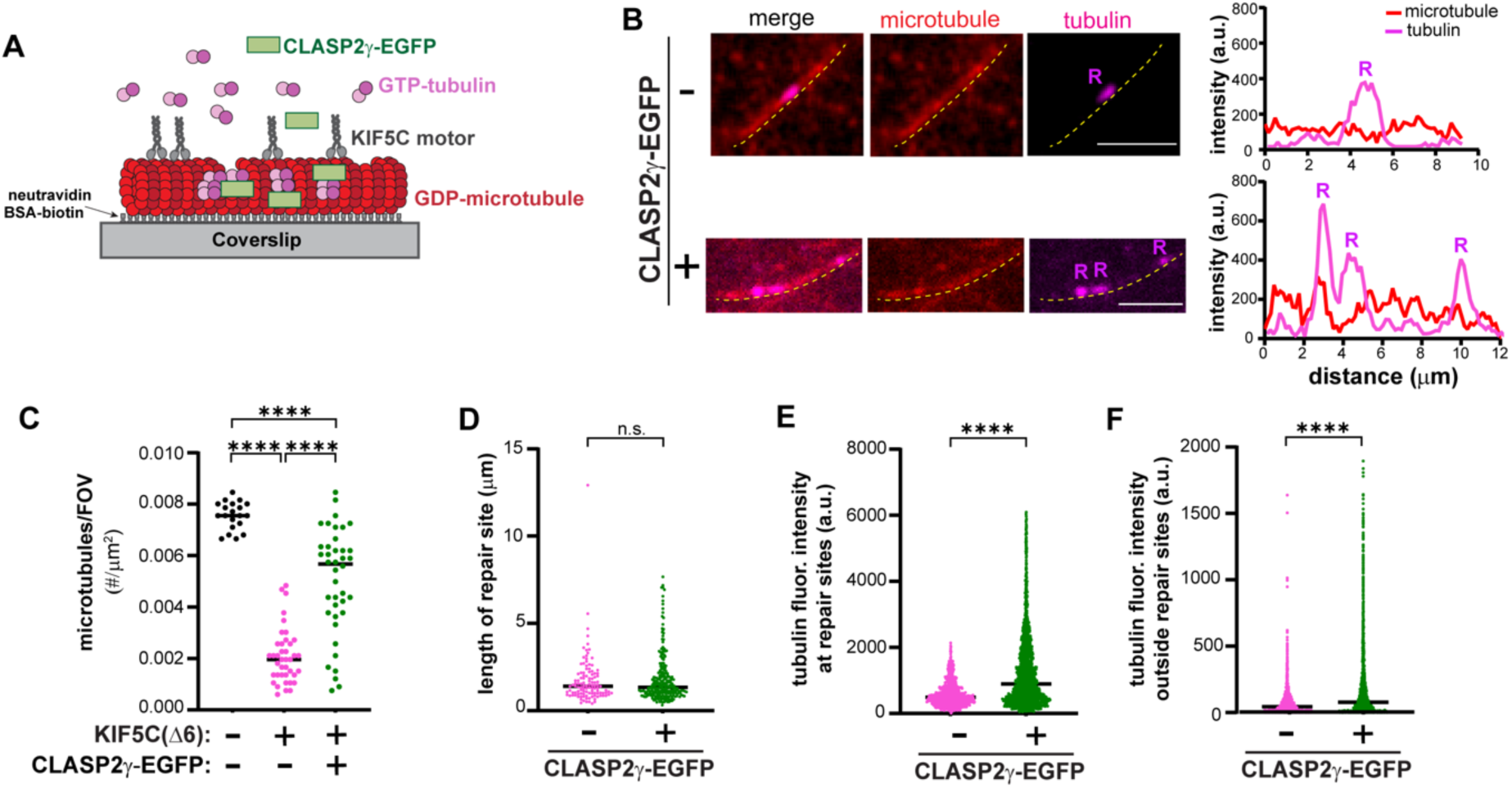
CLASP2γ promotes repair of motor-induced microtubule damage. (A) Schematic of microtubule repair assay. GMPCPP-capped GDP-microtubules were grown from GMPCPP-seeds attached on a coverslip. Purified KIF5C(Δ6)-Halo-FLAG protein (12 nM) was added to the flow chamber in the presence of 10 µM free Hilyte647-tubulin and in the absence or presence of CLASP2γ-EGFP. Static images were obtained after 7 min of free tubulin incorporation into motor-induced damage sites. (B) (left) representative images and (right) fluorescence intensity profiles of free Hilyte647-tubulin (magenta) incorporation into GDP-microtubules (red) in the absence or presence of 200 nM CLASP2γ-EGFP. R = repair site. Dotted yellow lines trace the microtubule. Scale bar: 10 µm. (C) Number of microtubules remaining per field of view (FOV) in the absence of KIF5C(Δ6) [0.008 ± 0.0001 (mean ± SE)] for n = 20 FOV across two independent experiments, in the presence of KIF5C(Δ6) (0.002 ± 0.0002) for n = 40 FOV across four independent experiments, or in the presence of KIF5C(Δ6) and CLASP2γ-EGFP (0.005 ± 0.0003) for n= 40 FOV across four independent experiments. ****p <0.0001 (Welch’s t test). (D) Quantification of the number and length of lattice repair sites. N = 106 repair sites with median length of 1.40 µm in the absence of CLASP2γ and n = 263 repair sites with median length of 1.33 µm in the presence of 200 nM CLASP2γ across four independent experiments. n.s.: not significant (Mann Whitney). (E) Quantification of fluorescence intensity of Hilyte647-tubulin incorporated at lattice repair sites in the absence or presence of 200 nM CLASP2γ-EGFP across four independent experiments. Median intensity = 492.30 a.u. in the absence of CLASP2γ-EGFP, and = 899.8 a.u. in the presence of 200 nM CLASP2γ-EGFP. ****p <0.0001 (Mann Whitney). (F) Quantification of the fluorescence intensity of Hilyte647-tubulin incorporated outside of the lattice repair sites. Median intensity = 43.5 a.u. in the absence of CLASP2γ-EGFP (n = 82 microtubules) and = 76.9 a.u. in the presence of 200 nM CLASP2γ-EGFP (n = 184 microtubules) across three independent experiments. ****p <0.0001 (Mann Whitney).

To determine whether CLASP2γ facilitates repair events, we quantified the incorporation of free tubulin into the lattice. We defined a lattice repair site as a bright punctum of new tubulin subunits incorporated into a pre-existing lattice. We found that although there was a 2.48-fold increase in the number of repair sites in the presence of CLASP2γ-EGFP (n = 263 repair sites) compared to the control (n = 106 repair sites), this increase scales with the increase in the number of microtubules remaining in the presence of CLASP2γ-EGFP (avg 0.005 ± 0.0003 microtubules/µm) compared to the control (avg 0.002 ± 0.0002 microtubules/µm) (Fig. 5C). We also found that there was no significant difference in the length of the repair sites in the presence of CLASP2γ-EGFP (median length 1.33 µm) compared to the control (median length 1.40 µm) (Fig. 5D). These results suggest that CLASP2γ does not prevent microtubule destruction by increasing the number or size of repair sites.

We then quantified the fluorescence intensity of the free tubulin incorporated along the microtubule lattice. We found that the tubulin intensity at repair sites was significantly increased in presence of CLASP2γ-EGFP (Fig. 5E). Interestingly, we found that the tubulin intensity outside of the repair sites also increased in presence of CLASP2γ-EGFP (Fig. 5F). These results indicate that CLASP2γ protects damaged microtubules from destruction by enhanced incorporation of free tubulin into repair sites. Furthermore, CLASP2γ appears to facilitate the incorporation of free tubulin into small damage sites and thereby prevent them from becoming large damage sites.

Finally, we verified that the ability of CLASP2γ to prevent microtubule destruction requires the incorporation of free tubulin into a KIF5C(Δ6)-damaged lattice. We performed lattice repair assays in the absence of free tubulin and found that under these conditions, the addition of CLASP2γ-EGFP was unable to prevent microtubule destruction (Fig. S5). This observation provides further support for the conclusion that CLASP2γ protects microtubules by promoting repair through the incorporation of free tubulin.

## Discussion

Recent work has revealed that the microtubule lattice is a plastic medium and that the stepping of molecular motor proteins along the microtubule surface can induce conformational changes in tubulin subunits and create damage sites [6, 8]. How cells repair lattice damage is a key question in the field. In this work, we investigate the role of EB1 and CLASP2γ in repairing lattice damage induced by the stepping of kinesin-1. Our results demonstrate that CLASP2γ is sufficient to protect microtubules from kinesin-induced damage.

To unravel the cellular mechanisms underlying microtubule repair, we used a targeted siRNA screen to investigate the role of specific MAPs in modulating microtubule damage in cells expressing either KIF5C(WT) or KIF5C(Δ6). Our goal was to identify proteins whose loss of expression leads to even more microtubule damage and destruction than induced by KIF5C alone. There are several limitations to the screen as performed. For example, we were unable to confirm the levels of knockdown for each siRNA target. In addition, the low number of cells analyzed for some conditions precludes making strong conclusions. Nevertheless, the screen identified EB1, DCTN1, and CLASP2 proteins as potential factors involved in repair of kinesin-induced lattice damage. In the future, a larger-scale screening approach could be employed to investigate additional cellular factors involved in lattice repair and structural integrity.

Interestingly, siRNAs targeting several MAPs led to less microtubule destruction than induced by kinesin expression alone. For example, loss of expression of the TOG-containing proteins CKAP5 (ch-TOG), TOGORAM1 (crescerin-1), TOGORAM2 (crescerin-2), or GCN1 resulted in less microtubule breakage and fragmentation in cells expressing KIF5C(WT) or KIF5C(Δ6). CKAP5 and the TOGORAM proteins belong to the MAP215 family proteins that bind to microtubule ends and promote polymerization whereas CLASP family proteins act as microtubule stabilizers by binding to the microtubule lattice to promote rescue and prevent depolymerization [41]. Further work is needed to understand the mechanism by which loss of CLASP family proteins leads to increased microtubule destruction whereas loss of MAP215 family proteins leads to reduced microtubule destruction.

While loss of EB proteins, particularly EB1, led to increased microtubule destruction in cells expressing KIF5C(WT) or KIF5C(Δ6), the addition of EB1 was not sufficient to protect microtubules in the *in vitro* microtubule destruction assay. This finding demonstrates that EB1 does not play a direct role in repairing kinesin-induced lattice damage. This finding is consistent with previous work demonstrating that EB1 is recruited after incorporation of GTP-tubulin into the lattice. Specifically, both EB1 and EB3 localize to repair sites and their localization is dependent on the incorporation of free GTP-tubulin at the damage site [23–25]. EB1 may thus play a role in amplifying the repair process in cells. In this scenario, the initial incorporation of GTP-tubulin would recruit EB proteins which can then act as a scaffold for recruitment of other plus-end tracking proteins, including CAP-Gly and TOG domain-containing factors which are implicated in microtubule stabilization and repair. Loss of EB1 in cells may thereby reduce repair efficiency and increase susceptibility to kinesin-induced damage.

In contrast to EB1, we could confirm a role for CLASP2 in the repair of KIF5C-induced damage both in cells and *in vitro*. In cells, a decrease in CLASP2 expression resulted in increased microtubule destruction, suggesting that CLASP2 is necessary for microtubule repair. *In vitro*, the addition of CLASP2γ-EGFP resulted in an increased number of microtubules remaining over time in both the microtubule destruction assay and the lattice repair assay, indicating that CLASP2 is sufficient for lattice repair.

We found that the addition of CLASP2γ-EGFP resulted in an increased amount of free tubulin incorporated at damage sites in the lattice repair assay. This result supports the hypothesis that CLASP2 directly facilitates the repair of kinesin-1-induced lattice damage. This result is consistent with the results of Aher et al [39] who found that the addition of GFP-CLASP2α results in a higher tubulin intensity at laser-induced damage sites and an increased frequency of complete lattice repair upon Taxol washout.

In addition, we found that there was an increase in tubulin intensity along the lattice (outside of repair sites) in the presence of CLASP2γ-EGFP. This finding suggests that CLASP2γ stabilizes microtubules not only by facilitating complete repair of large lattice defects, but also by repairing small damage sites generated during kinesin stepping (Fig. 5). Presumably, the small tubulin incorporation sites are thereby prevented from progressing into larger lattice damage sites that lead to microtubule breakage and depolymerization. These small repair sites were not observed in previous assays in which damage was mechanically induced at a single site along the microtubule [39].

In conclusion, our results demonstrate that CLASP2γ has a direct role in promoting microtubule repair in response to motor-induced damage and preventing catastrophic microtubule disassembly. Our findings are consistent with previous work demonstrating that CLASPs are necessary for stabilization of microtubules in response to mechanical forces [39, 49, 50]. We propose that CLASP2 plays a protective role by stabilizing exposed GDP-tubulins at damage sites, similar to how CLASPs stabilize a pre-catastrophe intermediate state at microtubule ends to suppress microtubule catastrophe [42, 46]. The CLASP family of TOG domain-containing MAPs thus appear to play a critical role in recognition, stabilization, and repair of vulnerable sites within the microtubule lattice as well as at the ends of microtubules.

## Key resources tables

**Table 1:** siRNA screen targets.

| Gene Symbol | Protein Name | UniProt ID |
| --- | --- | --- |
| MAPRE1 | Microtubule-associated protein RP/EB family member 1 (EB1) | Q15691 |
| MAPRE2 | Microtubule-associated protein RP/EB family member 2 (EB2) | Q15555 |
| MAPRE3 | Microtubule-associated protein RP/EB family member 3 (EB3) | Q9UPY8 |
| CLIP1 | CAP-Gly domain containing linker protein 1 (CLIP-170) | P30622 |
| CLIP2 | CAP-Gly domain containing linker protein 2 (CLIP-115) | Q9UDT6 |
| CLIP3 | CAP-Gly domain containing linker protein 3 (CLIPR-59) | Q96DZ5 |
| CLIP4 | CAP-Gly domain-containing linker protein 4 (CLIP4) | Q8N3C7 |
| DCTN1 | Dynactin subunit 1 (DCTN1, also known as p150-glued) | Q14203 |
| TBCB | Tubulin-folding cofactor B (TBCB, also known as CKAP1) | Q99426 |
| TBCE | Tubulin-folding cofactor E (TBCE) | Q15813 |
| CLASP1 | cytoplasmic linker associated protein 1 (CLASP1) | Q7Z460 |
| CLASP2 | cytoplasmic linker associated protein 2 (CLASP2) | O75122 |
| CKAP5 | Cytoskeleton-associated protein 5 (CKAP5, also known as ch-TOG or MAP215) | Q14008 |
| TOGARAM2 | TOG array regulator of axonemal microtubules 2 (TOGARAM2, also known as crescerin-2 or FAM179A) | Q6ZUX3 |
| TOGARAM1 | TOG array regulator of axonemal microtubules 1 (TOGARAM1, also known as crescerin-1 or FAM179B) | Q9Y4F4 |
| GCN1 | GCN1 eIF-2-alpha kinase activator homolog | E1NZA1 |

**Table 2:**
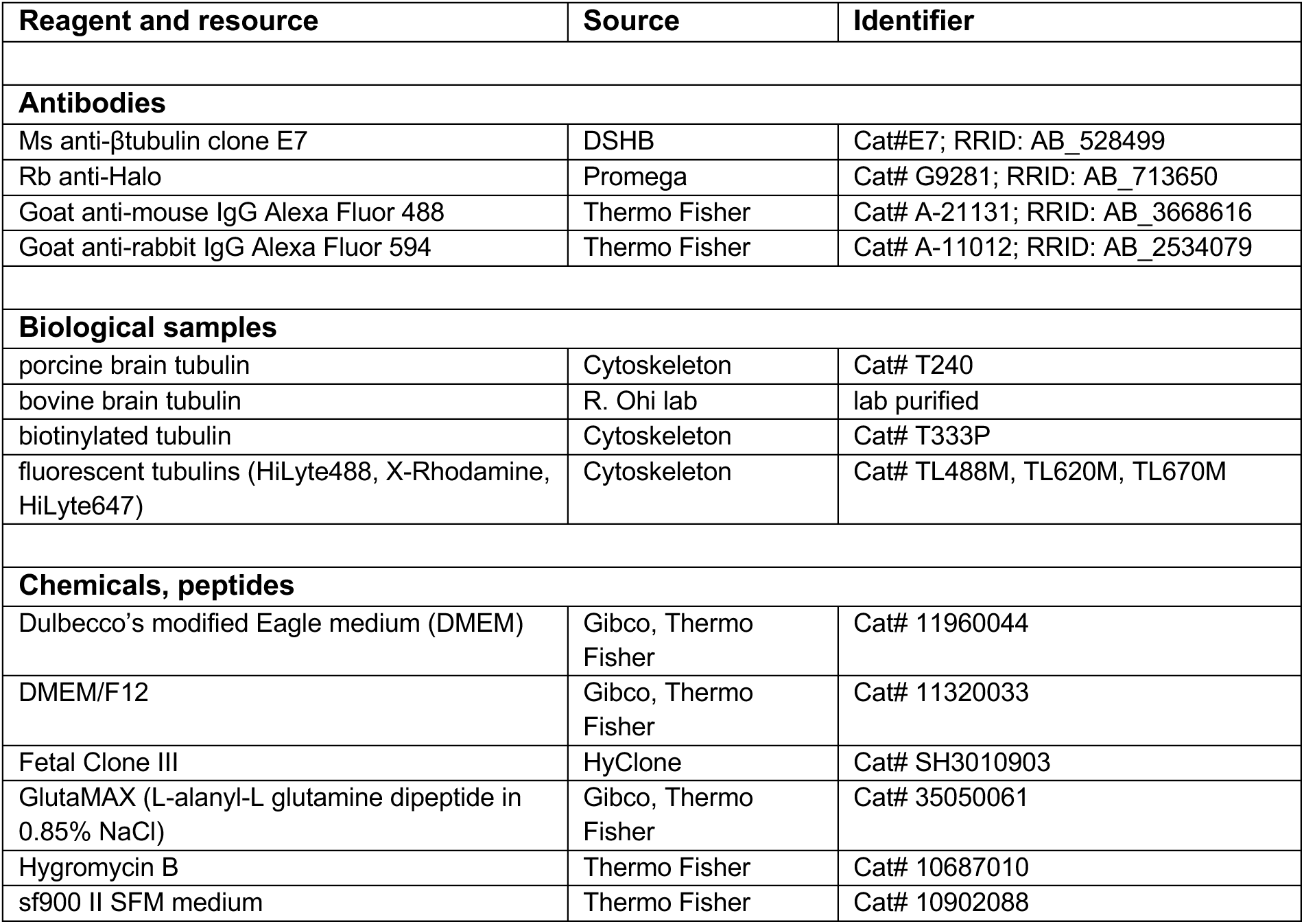
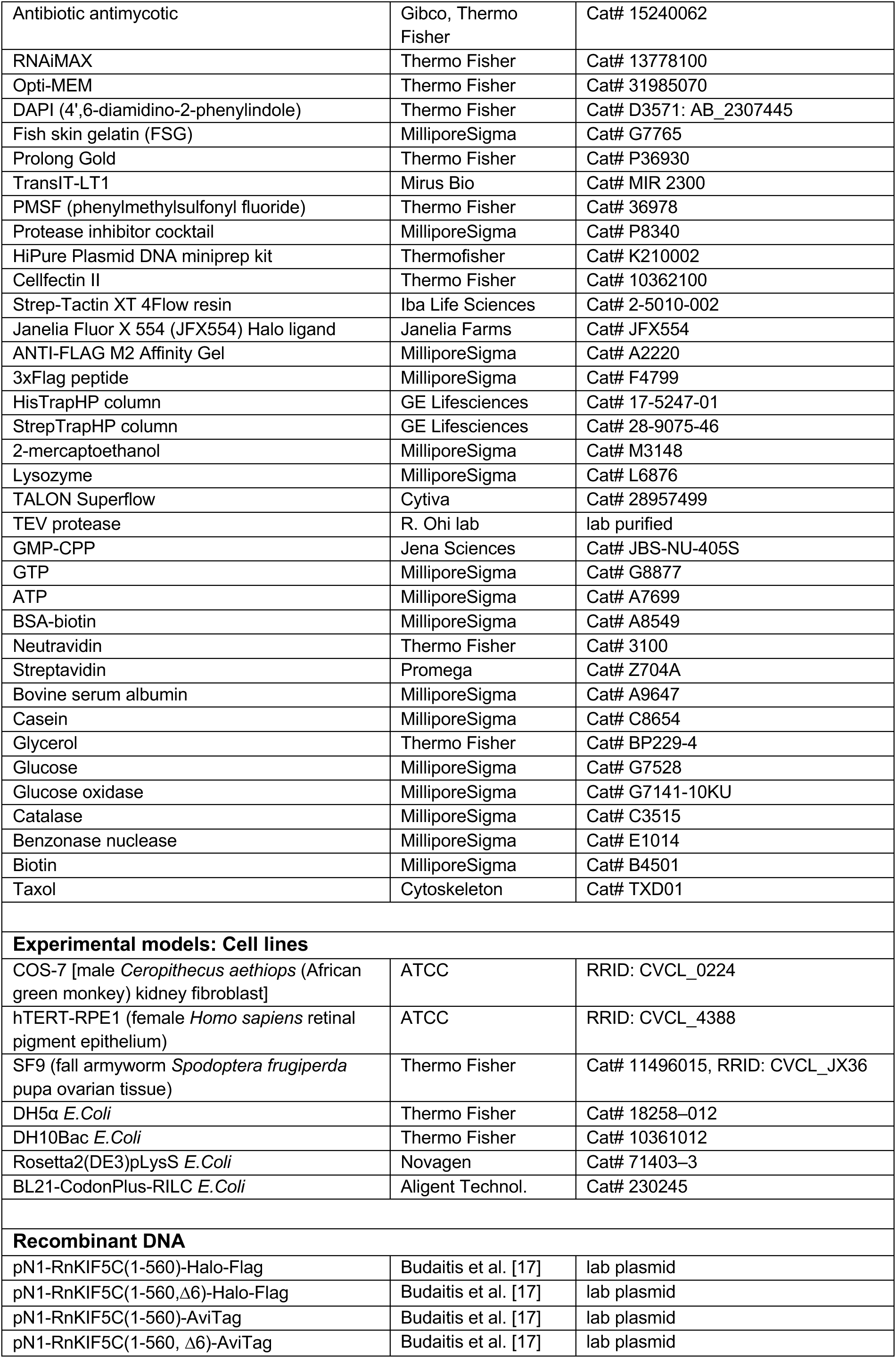
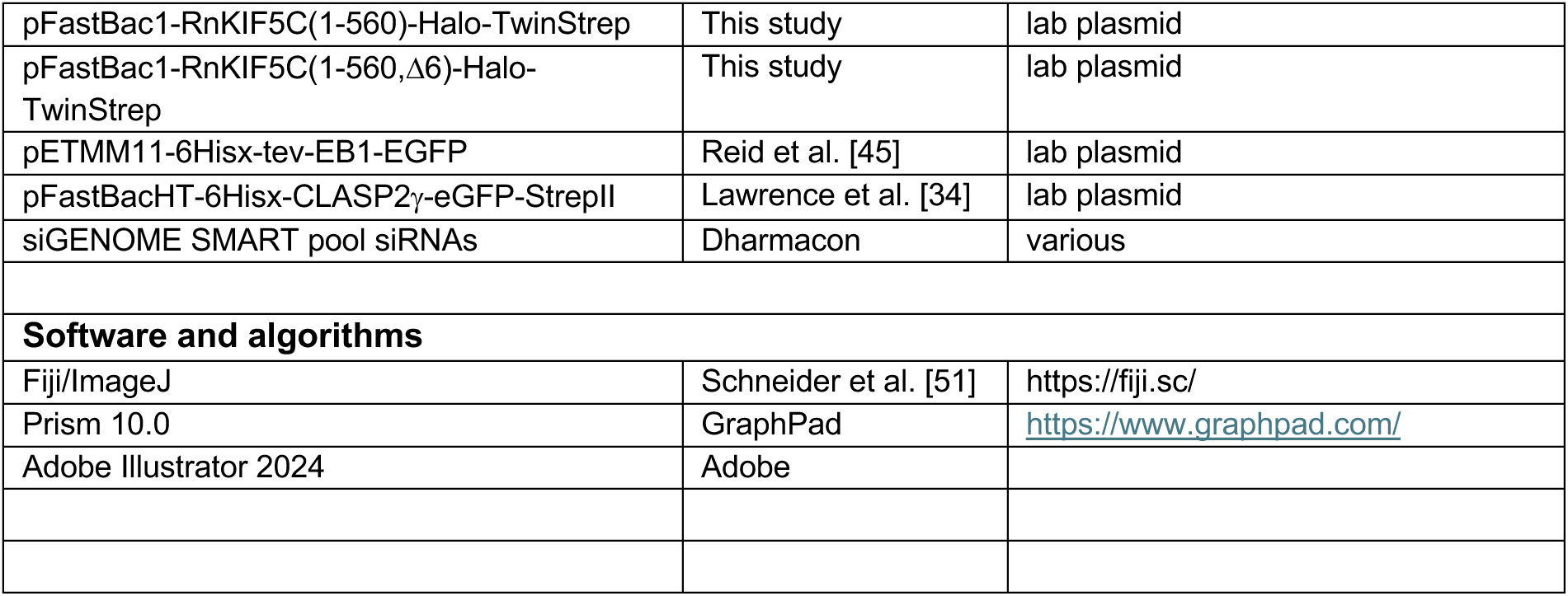
Reagents.

## Methods

### Cell culture

COS-7 cells were grown in Dulbecco’s Modified Eagle Medium (DMEM) supplemented with 10% (vol/vol) Fetal Clone III and 2 mM GlutaMAX at 37 °C with 5% CO_2_. hTERT-RPE1 cells were grown in DMEM/F12 (1:1) with 10% (vol/vol) Fetal Clone III, 0.5 mg/ml hygromycin B, and 2 mM GlutaMAX at 37 °C with 5% CO_2_. Sf9 cells were cultured in suspension with serum-free sf900 II SFM medium supplemented with antibiotic antimycotic at 28°C in a non-CO_2_ non-humidified incubator with an orbital shaker platform set at 110 rpm. All cell lines are checked annually for mycoplasma contamination and COS-7 cells were authenticated through mass spectrometry (the protein sequences exactly match those in the *Ceropithecus aethiops* genome).

### Plasmids

A truncated, constitutively active kinesin-1 [rat KIF5C(1-560)] plasmid was used. The Δ6 version lacks aa 2-6 and is described in [17]. KIF5C(1-560) was tagged with Halo-Flag tags for immunofluorescence, single-molecule imaging assays, and microtubule repair assays. For microtubule destruction assays KIF5C(1-560) was tagged with Halo-TwinStrep tags or was biotinylated via an AviTag (aa sequence GLNDIFEAQKIEWHE) and co-expression with HA-BirA as described [52]. EB1-EGFP protein was expressed and purified using plasmid pETEMM1-6xHis-tev-EB1-EGFP [45] and CLASP2γ-EGFP-StrepII protein was expressed and purified using plasmid pHAT-6xHis-CLASP2γE-StrepII [34]. All plasmids were verified by DNA sequencing.

### siRNA knockdown, transfection, and immunofluorescence in hTERT-RPE1 cells

siGENOME SMARTpool siRNAs were mixed with RNAiMAX in OptiMEM and incubated for 15 min at room temperature. Coverslips were placed in each well of a 12-well plate and then the siRNA mix was added followed by 1×10^5^ hTERT-RPE1 cells in DMEM:F12/FCIII/GlutaMAX. After 4-6 hr, the media was replaced with fresh DMEM:F12/FCIII/hygro/GlutaMAX. 24 hr later the cells were transfected with plasmids for expression of KIF5C(1-560)-Halo-Flag or KIF5C(1-560,Δ6)-Halo-Flag using Trans-IT LT1 and Opti-MEM. 24 hr later the cells were subjected to a second round of siRNA transfection. 24 hr later the cells were rinsed with PBS^+^ (PBS with 0.9 mM CaCl_2_, 0.5 mM MgCl_2_), fixed with 3.7% paraformaldehyde in PBS^+^, permeabilized with 0.2% Triton X-100 in PBS^+^, blocked with 0.2% fish skin gelatin (FSG) in PBS^+^, and then stained with primary and secondary antibodies in blocking buffer. Primary antibodies were Ms anti-β-tubulin clone E7 (1:1000) and Rb anti-Halo (1:1000). Secondary antibodies were AlexaFluor488 anti-Ms (1:500) and AlexaFluor594 anti-Rb (1:500) with 14.3 µM DAPI. After washing with FSG/PBS^+^ and PBS^+^, the cells were mounted on Prolong Gold on a glass microscope slide. Images were acquired on an inverted epifluorescence microscope (Nikon TE200E) with a 60x, 1.4 NA oil-immersion objective and an Orca Flash4 OLT digital CMOS camera (Hamamatsu).

### Transfection and preparation of COS-7 cell lysates containing biotinylated KIF5C-AviTag motors

COS-7 cells were transfected with plasmids for expressing HA-BirA with KIF5C(1-560)-AviTag or KIF5C(1-560,τι6)-AviTag using TransIT-LT1 and Opti-MEM. BirA efficiently biotinylates the AviTag in cells. 24 hr later the cells were trypsinized and harvested by centrifugation at 3000 x g at 4°C for 3 min. The cell pellet was resuspended in cold 1X PBS, centrifuged at 3000 x g at 4°C for 3 min, and the pellet was resuspended in 25 µL of cold lysis buffer [25 mM HEPES/KOH, 115 mM potassium acetate, 5 mM sodium acetate, 5 mM MgCl_2_, 0.5 mM EGTA, and 1% (vol/vol) Triton X-100, pH 7.4] with 1 mM ATP, 1 mM PMSF, and 1% (vol/vol) protease inhibitor cocktail. Lysates were clarified by centrifugation at 20,000 x g at 4°C for 10 min and lysates were snap frozen in 5 µL aliquots in liquid nitrogen and stored at - 80°C.

### Protein expression and purification

#### KIF5C motor protein from SF9 cells

Plasmids pFastBac1-KIF5C(1-560)-Halo-TwinStrep or pFastBac1-KIF5C(1-560,τι6)-Halo-TwinStrep were transformed into DH10Bac *E. coli* to generate recombinant bacmids. Bacmid DNA was isolated with the HiPure Plasmid DNA miniprep kit, confirmed by PCR analysis, and transfected into Sf9 cells using Cellfectin II. 7d after transfection, the supernatant containing P1 baculovirus was collected by centrifugation at 3,000 rpm for 3 min at 4°C. The baculovirus was amplified by successive infection of Sf9 cells to generate P2 and P3 baculoviruses. Baculovirus-containing supernatants were stored at 4°C in the dark. For protein production and purification, Sf9 cells were infected with 3% (vol/vol) P3 baculovirus. 3 d after infection, the cells were harvested by centrifugation for 15 min at 3,000 rpm at 4°C. The pellet was washed once with PBS and resuspended in ice-cold lysis buffer (200 mM NaCl, 4 mM MgCl_2_, 0.5 mM EDTA, 1 mM EGTA, 0.5% igepal, 7% sucrose, and 20 mM imidazole-HCl, pH 7.5) supplemented with 2 mM ATP, 1 mM PMSF, 5 mM DTT, and protease inhibitor cocktail. After 30 min incubation on ice, the lysates were clarified by ultracentrifugation for 20 min at 20,000 rpm in F12-8×50y rotor (Sorvall 3421). The supernatant was incubated with Strep-Tactin XT 4Flow beads for 1hr at 4°C with rotation. The beads were drained in a PD-10 column and washed with wash buffer (150 mM KCl, 25 mM imidazole-HCl, pH 7.5, 5 mM MgCl_2_, 1 mM EDTA, and 1 mM EGTA) supplemented with 1 mM PMSF, 3 mM DTT, 3 mM ATP, and protease inhibitor cocktail. Bound proteins were eluted with elution buffer (25 mM KCl, 25 mM imidazole-HCl, pH 7.5, 5 mM EGTA, 2 mM MgCl_2_, 2 mM DTT, 0.1 mM ATP, 1 mM PMSF, protease inhibitor cocktail and 10% glycerol) supplemented with 50 mM biotin in 6×0.5 mL fractions. SDS-PAGE of the eluted fractions was done, and pure fractions were selected by looking at the bands on a Coomassie-stained gel. Pure protein fractions were combined and dialyzed in dialysis buffer (25 mM imidazole-HCl, pH 7.5, 25 mM KCl, 5 mM EGTA, 2 mM MgCl_2_, 2 mM DTT, 0.1 mM ATP and 10% glycerol) at 4°C to remove biotin from the sample. After 2 hrs, the buffer was changed with fresh dialysis buffer and dialyzed overnight at 4°C. The protein sample was collected by centrifugation, and aliquots were snap frozen in liquid nitrogen and stored at -80°C.

#### KIF5C(1-560,Δ6) motor protein from mammalian cells

COS-7 cells were transfected with plasmids for expression of KIF5C(1-560,Δ6)-Halo-Flag and the protein was fluorescently labelled by the inclusion of 50 nM JFX554 Halo ligand in the growth medium. Cells from two 10 cm dishes were harvested 24 hr after transfection and lysed in 1 ml lysis buffer [25 mM HEPES, 115 mM KOAc, 5 mM NaOAc, 5 mM MgCl_2_, 0.5 mM EGTA, 1% Triton X-100, pH to 7.4 with KOH] supplemented with protease inhibitor cocktail, 1 mM PMSF, 1 mM ATP, and 1 mM DTT. After centrifugation at 16,000xg for 10 min at 4 C, the supernatant was incubated with 50 µl anti-Flag M2 agarose beads with rotation for 1.5 hr at 4°C. The beads were washed with wash buffer (150 mM KCl, 20 mM Imidazole pH 7.5, 5 mM MgCl_2_, 1 mM EDTA, 1 mM EGTA) supplemented with protease inhibitor cocktail, 1 mM PMSF, 1 mM DTT, and 3 mM ATP, and washed again with wash buffer supplemented with protease inhibitor cocktail, 1 mM PMSF and 1 mM DTT. The protein was eluted with 80 µl BRB80 buffer (80 mM PIPES/KOH pH 6.8, 1 mM MgCl_2_, 1 mM EGTA) supplemented with protease inhibitor cocktail, 1mM PMSF, 0.5mM DTT, 0.1mM ATP, 0.5 mg/ml 3xFlag peptide for 1hr. The protein was collected as the supernatant after centrifugation at 1,500xg for 5 min at 4°C. Aliquots were snap-frozen in liquid nitrogen and stored at -80°C.

#### CLASP2γ-EGFP protein from SF9 cells

His-CLASP2γ-eGFP-StrepII protein was expressed in and purified from baculovirus-infected Sf9 insect cells using the Bac-to-Bac system as described previously [34]. Briefly, a modified pHAT vector for insect cell expression containing 6xHis-CLASP2γ-eGFP-StrepII was transformed into DH10Bac cells and bacmid DNA was purified using the HiPure Plasmid DNA miniprep. Bacmid DNA was transfected into Sf9 calls and baculovirus-infected insect cells (BIIC) stocks were prepared as previously described [53]. Sf9 cells were infected at a density of 1 × 10^6^ viable cells/ml with BIIC stocks at a ratio of 10^−4^ BIIC:total culture volume. Cells were harvested 5 d after infection. Cell pellets were lysed by one freeze–thaw cycle and Dounce homogenizing in lysis buffer [50 mM HEPES (4-(2-hydroxyethyl)- 1-piperazineethanesulfonic acid) (pH 7.5), 120 mM KCl, 5% glycerol, 0.1% Tween-20, 2 mM MgCl_2_, 10 mM imidazole, and 1 mM DTT] containing protease inhibitors. Crude lysates were clarified by centrifugation for 20 min at 4°C at 35,000 rpm in a Beckman L90K Optima centrifuge and 50.2 Ti rotor. Clarified lysates were applied to a HisTrapHP column according to the manufacturer’s protocol. His-tagged proteins were eluted with elution buffers [50 mM HEPES (pH 7.5), 120 mM KCl, 5% glycerol, 0.1% Tween-20, 2 mM MgCl_2_, 1 mM DTT, and 300 mM imidazole]. Peak His-CLASP2γ-eGFP-StrepII elution fractions were pooled and applied to a StrepTrapHP column according to the manufacturer’s protocol. Purified His-CLASP2γ-eGFP-StrepII was eluted with 2.5 mM desthiobiotin. Aliquots were snap-frozen in liquid nitrogen and stored at -80°C.

#### EB1-EGFP protein from bacteria cells

EB1-EGFP was purified as described previously [45]. Briefly, plasmid pETMM11-6xHis-tev-EB1-EGFP was transformed into Rosetta (DE3) pLysS *E. coli*, grown in 10 ml of LB+kanamycin at 37° C to an A600 of 0.44. IPTG was then added to 2 mM and the culture was incubated with shaking at 16° C for 20 hrs. The culture was centrifuged at 4000 rpm for 45 min at 4°C. The cell pellet was resuspended with 5 times of weight of the pellet in PBS, 0.1% tween-20, 5 mM 2-mercaptoethanol, 1 mg/ml lysozyme and 1x protease inhibitors. The cells were mixed at 4° C for 2 hr and then sonicated on ice at 70% power, 50% duty, 6 min. The lysate was centrifuged at 20,000 rpm at 4° C. The supernatant was passed through 2 ml of Talon SuperFlow Resin. The resin was sequentially washed with five column volumes Buffer A (50 mM sodium phosphate pH7.5, 300 mM KCl, 10% glycerol, 5 mM 2-mercaptoethanol), followed by 95% buffer A, 5% buffer B (Buffer A/300 mM imidazole), followed by 90% A/10% B, and finally 85% Buffer A/15% Buffer B. All buffers contained 1x protease inhibitors cocktail. Protein was eluted with 1 ml fractions of 100% Buffer B + 1xPI cocktail until the eluted fractions look green. Elution fractions were analyzed on SDS PAGE with Coomassie staining. Relevant fractions were combined and eluted through desalting columns with buffer A. The eluted protein was digested with 6xHIS-tagged TEV protease at a 1:10 w/w protein ratio overnight at 4° C. TEV-cleaved protein solution was incubated with 100 µL talon resin for 60 min at 4°C to remove cleaved 6xHIS tag. After centrifugation at 500xg, 5 min at 4°C, the supernatant was collected containing EB1-EGFP. The EB1-EGFP solution was concentrated, and aliquots were snap-frozen in liquid nitrogen and stored at -80°C.

### Total Internal Reflection Fluorescence (TIRF) assays

#### Preparation of glycerol-stabilized microtubules

A tubulin mixture was prepared containing 20 mg/mL unlabeled bovine brain tubulin (gift of R. Ohi, University of Michigan) and 2.5 mg/ml Hilyte 647-tubulin (Cytoskeleton) in 100 mM DTT, 100 mM MgCl_2_, 25 mM GTP. The tubulin mix was incubated at 37°C for 35 min. Prewarmed 25% glycerol in BRB80 (80 mM PIPES/KOH pH 6.8, 1 mM MgCl_2_, 1 mM EGTA) containing 1 mM GTP was added to the polymerized microtubule solution and centrifuged at 15,000 rpm for 15 min at room temperature. The supernatant was aspirated and the pellet was resuspended with 25% glycerol in BRB80 containing 1 mM GTP. The resulting glycerol-stabilized GDP-microtubule solution was maintained at 37°C and used within 2-4 hr after polymerization.

#### Microtubule destruction assay

KIF5C(1-560)-AviTag motors in cell lysates or purified KIF5C(1-560)-Halo-TwinStrep motors were attached to the surface of a coverslip by sequential incubation (5 min each) of a flow chamber with (i) 1 mg/ml BSA-biotin, (ii) wash buffer (0.5 mg/ml casein in BRB80), (iii) 0.5 mg/ml neutravidin for AviTag motors/or 1.0 mg/ml streptavidin for TwinStrep motors, (iv) wash buffer, (v) ∼200 nM AviTag motors in cell lysates or ∼400 nM purified TwinStrep motors, (vi) wash buffer, (vii) glycerol-stabilized GDP-microtubules, and (viii) wash buffer [1 mg/ml casein, 25% glycerol in P12 buffer (12 mM PIPES/KOH pH 6.8, 1 mM MgCl_2_, 1 mM EGTA)]. To initiate microtubule gliding by the surface-attached kinesin motors, motility mix was added to the chamber [1 mM DTT, 1 mM MgCl_2_, 10 mM glucose, 0.08 mg/mL catalase, 0.2 mg/mL glucose oxidase, 25% glycerol, supplemented with 2 mM ATP and 7 µM unlabeled tubulin (gift of R. Ohi, University of Michigan) in P12 buffer]. Images of microtubules in 9 fields of view were obtained every 5 min (0,5,10,15,20 min time points) using a Nikon Ti-E/B inverted TIRF microscope equipped with a 100x 1.49 N.A. oil immersion TIRF objective, three 20 mW diode lasers (488 nm, 561 nm and 640 nm), and EMCCD detector (Andor iXonX3DU897). The total number of microtubules in each field of view was manually counted using Fiji/ImageJ2, with only microtubules ≥ 2 µm in length included in the analysis. The measurements were averaged across 27 fields of view from three independent experiments for each time point. To quantify the length of microtubules, Ridge Detection plugin from Fiji/ImageJ2 was used in selected ROIs.

#### Single-molecule motility assays

Microtubules were polymerized from unlabeled and HiLyte-647-labeled porcine brain tubulin in BRB80 buffer (80 mM PIPES/KOH pH 6.8, 1 mM MgCl_2_, 1 mM EGTA) supplemented with 2 mM GTP and 2 mM MgCl_2_ and incubated for 60 min at 37 °C. Taxol in prewarmed BRB80 buffer was added to 2 mM final concentration and the microtubules were incubated for an additional 60 min. The microtubules were stored in the dark at room temperature for up to 2 weeks. Microtubules were diluted in fresh BRB80 buffer supplemented with 10 µM Taxol, infused into flow cells, and incubated for five minutes to allow for nonspecific absorption to the glass. Flow cells were incubated with (i) blocking buffer [30 mg/mL casein in P12 buffer (12 mM PIPES/KOH pH 6.8, 1 mM MgCl_2_, 1 mM EGTA) supplemented with 10 µM Taxol for four minutes and then (ii) motility mixture (100 pM KIF5C(1-560,Δ6)-Halo^554^-Flag, 25 mL P12 buffer, 15 mL blocking buffer, 1 mM ATP, 0.5 mL 100 mM DTT, 0.5 mL of 20 mg/mL glucose oxidase, 0.5 mL of 8 mg/mL catalase, and 0.5 mL 1M glucose) with 0 nM or 200 nM CLASP2γ-EGFP-StrepII. Flow chambers were sealed with molten paraffin wax and imaged on an inverted Nikon Ti-E/B TIRF microscope with a perfect focus system, a 100x 1.49 NA oil immersion TIRF objective, three 20 mW diode lasers (488 nm, 561 nm, and 640 nm) and EMCCD camera (Andor iXon+ DU879). Images were acquired every 50 ms for 30 s at room temperature. Acquisition was controlled using Nikon Elements software. Motility data were analyzed in Fiji/ImageJ by first generating maximum intensity projections to identify microtubule tracks and then generating kymographs along those microtubules (width = 3 pixels). Only motility events that lasted for at least three frames were analyzed. Furthermore, events that ended as a result of a motor reaching the end of a microtubule were included; therefore, the reported run lengths for highly processive motors are likely to be an underestimation. The velocities were binned, plotted as a histogram, fit to a Gaussian, and a two-tailed Welch’s t test was used to assess whether velocity distributions were significantly different between 0 nM and 200 nM CLASP2γ-EGFP-StrepII. The cumulative distributions of motor run lengths were done, and a Kolmogorow-Smirnov test was used to assess whether run length distributions were significantly different between 0 nM and 200 nM CLASP2γ-EGFP-StrepII conditions.

#### Microtubule repair assay

GMPCPP-microtubule seeds were prepared by polymerizing 25 µM tubulin (Cytoskeleton) consisting of 6% biotinylated-tubulin (Cytoskeleton)and 6% fluorescent (X-Rhodamine or HiLyte647) tubulin (Cytoskeleton) in the presence of the nonhydrolyzable GTP analogue GMPCPP (Jena Bioscience) in BRB80 buffer and 2.5 mM MgCl_2_ for 35 min at 37°C. The seeds were sedimented by centrifugation at 90,000 rpm for 5 min at 25°C (Beckman Coulter). The microtubule pellet was resuspended in warm BRB80 buffer and microtubule seeds were stored in the dark at room temperature. GMPCPP-seeds were attached to the surface of a coverslip by sequential incubation of a flow chamber with: (i) 1 mg/ml BSA-biotin for 5 min, (ii) blocking buffer, (iii) 0.5 mg/ml neutravidin for 5 min, (iv) blocking buffer, (v) GMPCPP-stabilized microtubule seeds (18% x-rhodamine tubulin) for 5 min, and (vi) blocking buffer. microtubules were polymerized from the seeds by incubating 26 µM tubulin [gift of R. Ohi (University of Michigan) with 12.5% x-rhodamine tubulin (Cytoskeleton)] and 1 mM GTP in imaging buffer [BRB80 buffer supplemented with 0.1% methylcellulose, 1 mg/ml casein, 3 mM MgCl_2_, 6 mM DTT and oxygen scavenger mix (16 mM glucose, 0.7 mg/ml catalase and 0.3 mg/ml glucose oxidase)] for 15 min. at 37°C. GDP-microtubules were capped by adding 13 µM unlabeled tubulin and 1mM GMPCPP in imaging buffer to the flow chamber for 5 min at 37°C. Wash buffer was flowed in to depolymerize uncapped microtubules. Subsequently, a mix containing 10 µM HiLyte647-tubulin, 1 mM GTP, 5 mM ATP, 12 nM purified KIF5C(1-560,Δ6)-Halo-Flag motors and 0 nM or 200 nM CLASP2γ-eGFP-StrepII in imaging buffer was flowed in. After incubation of motors with microtubules in the presence of free tubulin for 7 min at 37°C, the flow chamber was washed with blocking buffer to remove unincorporated HiLyte 647-tubulin and unbound motors and then 13 µM unlabeled tubulin was added to prevent microtubule depolymerization. The chamber was sealed with molten paraffin wax, and images were collected on a Nikon Ti-E/B TIRF microscope equipped with a 100X 1.49 N.A. oil immersion TIRF objective, three 20 mW diode lasers (488 nm, 561 nm and 640 nm), and EMCCD detector (Andor iXon X3DU897). Microtubule repair sites were defined as sites where HiLyte 647-tubulin incorporation into a microtubule was flanked on both sides by x-rhodamin tubulin lattice. The length of incorporation sites was quantified using Fiji/imageJ2. To quantify the intensity of HiLyte 647-tubulin incorporation in repair sites and outside the repair sites, pixel intensity was analyzed along the repair sites and subtracting the repair site pixel intensities from the whole microtubule pixel intensities respectively by Fiji/imageJ2. Background intensity was subtracted from all events by Fiji/imageJ2.

## ACKNOWLEDGEMENTS

This work was supported by grants R35GM131744 to KJV and R35GM1192552 to MZ from the National Institutes of Health. We thank Takashi Hotta (Ohi lab, University of Michigan) for advice on protein purification and Puck Ohi (Ohi lab, University of Michigan) for sharing purified tubulin. We thank Verhey lab members, Morgan DeSantis, Dave Sept, Mike Cianfrocco (all University of Michigan), and members of their laboratories for advice and feedback.

## AUTHOR CONTRIBUTIONS

**Jakia Jannat Keya**: Conceptualization; Investigation; Data curation; Formal analysis; Validation; Methodology; Writing—original draft; Writing—review and editing. **Rahul Riberio**: Investigation; Data curation; Formal analysis. **Yang Yue**: Methodology. **Elizabeth J. Lawrence**: Resources; Writing—review and editing. **Marija Zanic**: Resources; Supervision; Funding acquisition; Project administration; Writing—review and editing. **Kristen J Verhey**: Conceptualization; Resources; Supervision; Funding acquisition; Project administration; Writing—original draft; Writing—review and editing.

## AUTHOR NOTES

The authors declare no competing interests.

---

**Fig. S1.**
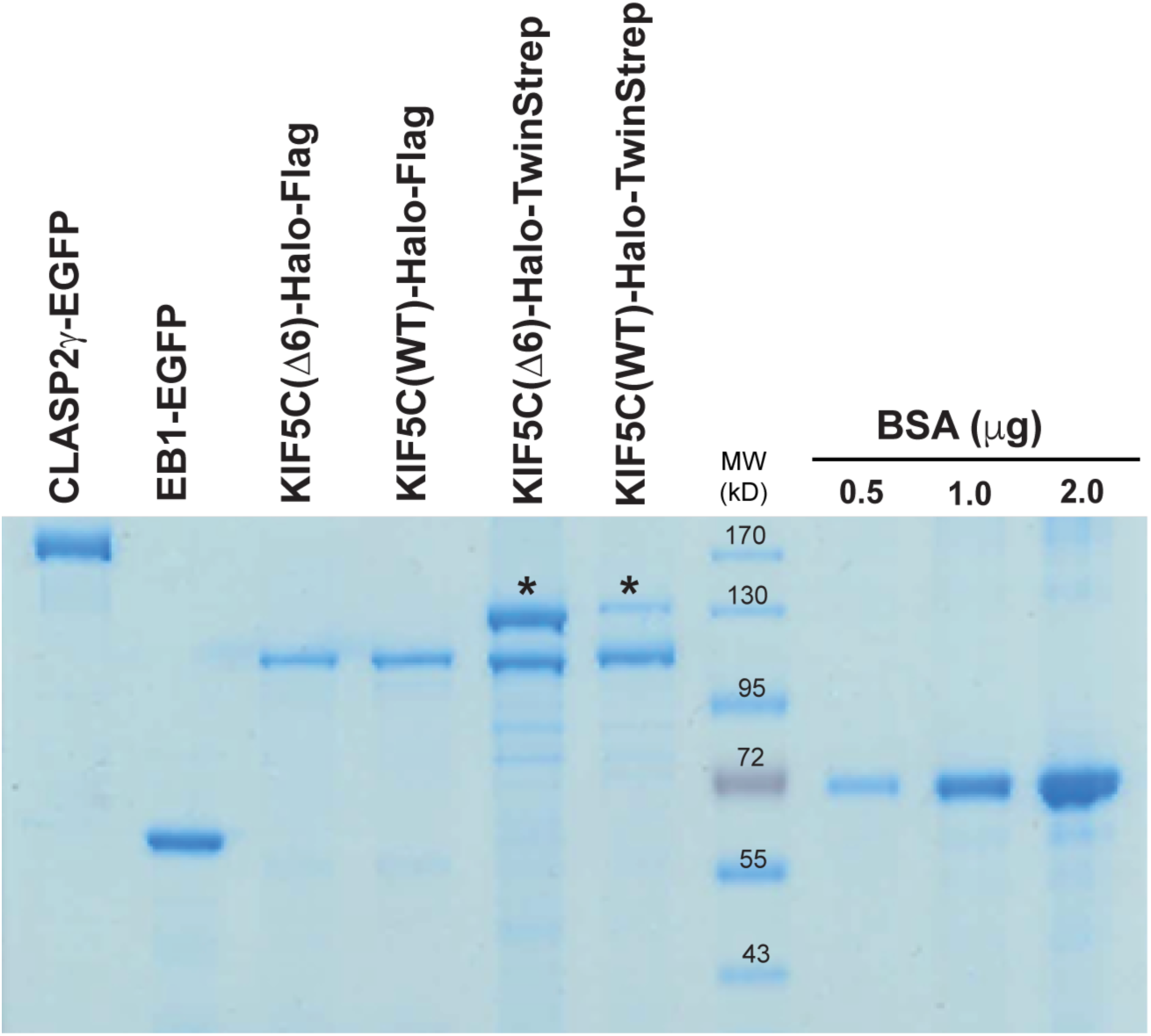
SDS-PAGE of purified proteins. Coomassie-stained gel of purified proteins used in this work and BSA standards. Asterisks indicate protein that co-purifies with KIF5C-Halo-TwinStrep and identified as pyruvate carboxylase by mass spectrometry. MW: molecular weight markers in kD.

**Fig S2.**
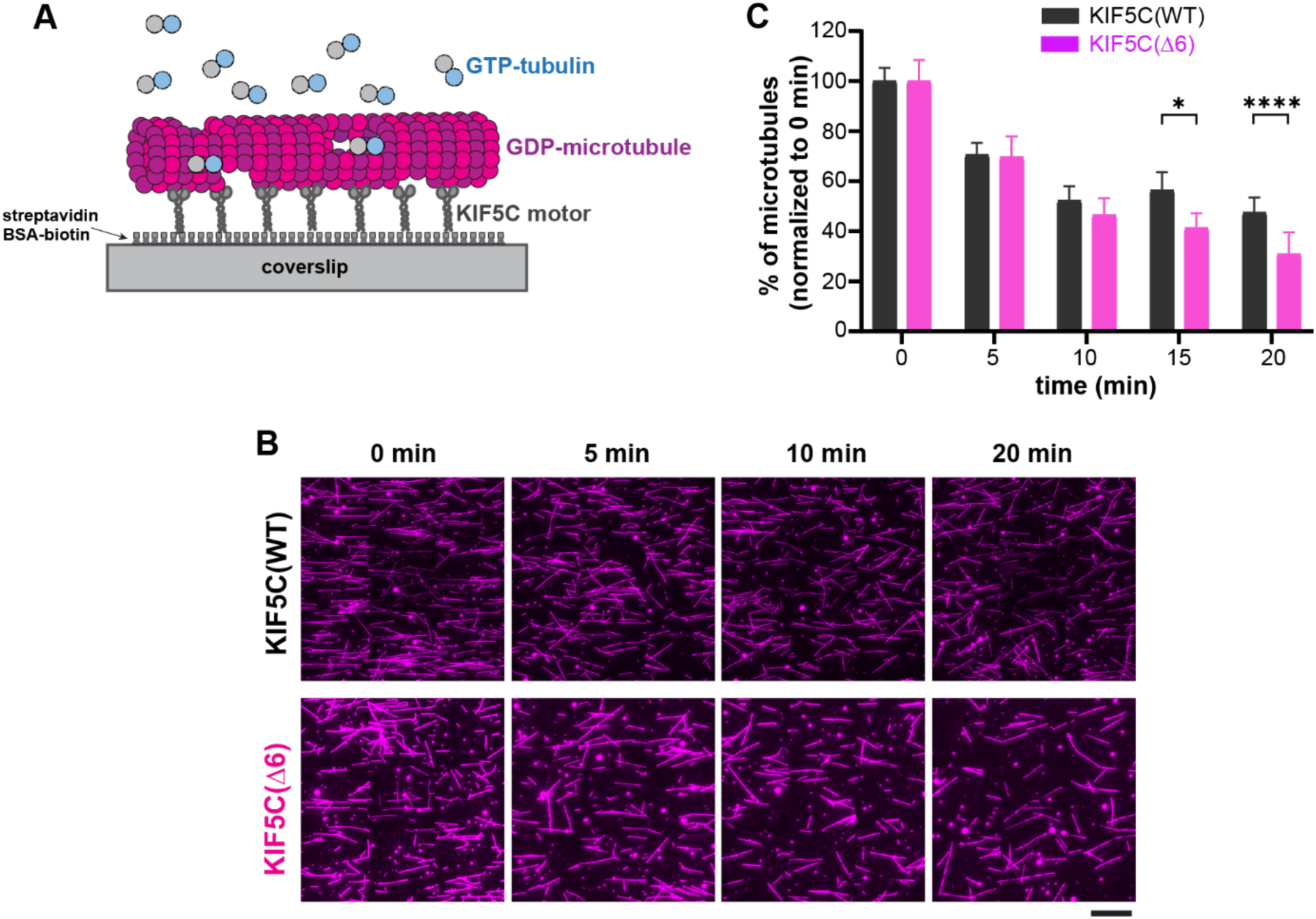
Microtubule destruction assay controls. (A) Schematic of microtubule destruction assay. KIF5C(WT) or KIF5C(Δ6) motors were attached to coverslips and then glycerol-stabilized GDP-microtubules were added in the presence of 2 mM ATP, 1 mM GTP and 7 µM unlabeled free tubulin. (B) Representative images of GDP-microtubules at 0, 5, 10, and 20 min after gliding driven by KIF5C(WT) or KIF5C(Δ6) motors. Scale bar: 20 µm. (C) Quantification of microtubule destruction over time by (black) KIF5C(WT) or (magenta) KIF5C(Δ6). The total number of microtubules was counted per field of view at 0 min (immediately after ATP addition to start KIF5C motility) and then after 5, 10, 15, or 20 min. The percent of microtubules remaining in each FOV was determined, averaged for 27 FOV across three independent experiments, and normalized to the 0 min time point (error bars: SE). * p<0.05, **** p<0.0001 (two-tailed t test).

**Fig S3.**
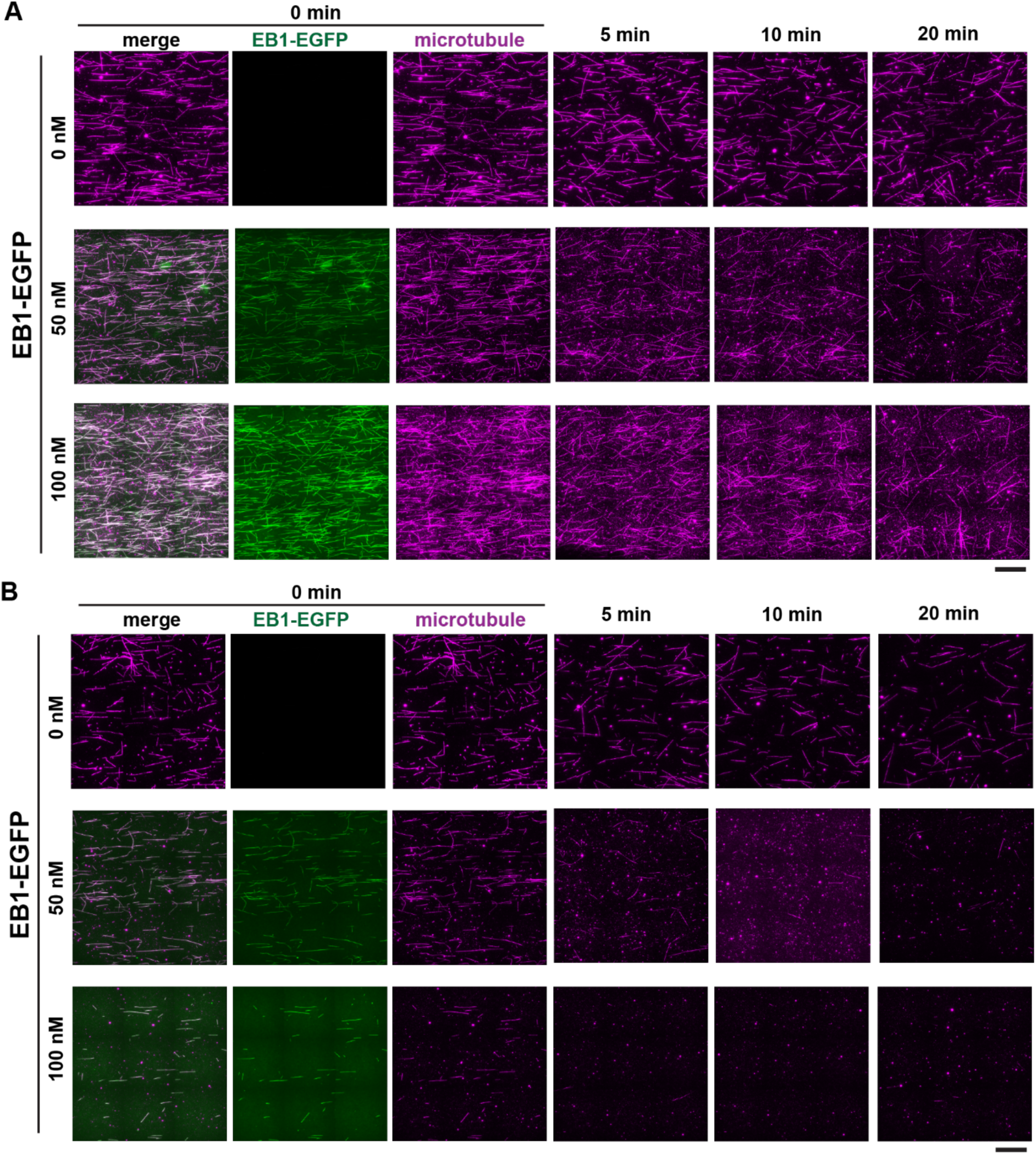
EB1 does not protect microtubules from kinesin-induced destruction. Representative images of microtubules remaining after damage induced by (A) KIF5C(WT) or (B) KIF5C(Δ6) in the absence or presence of the indicated concentrations of EB1-EGFP. Scale bar: 20 µm.

**Fig S4.**
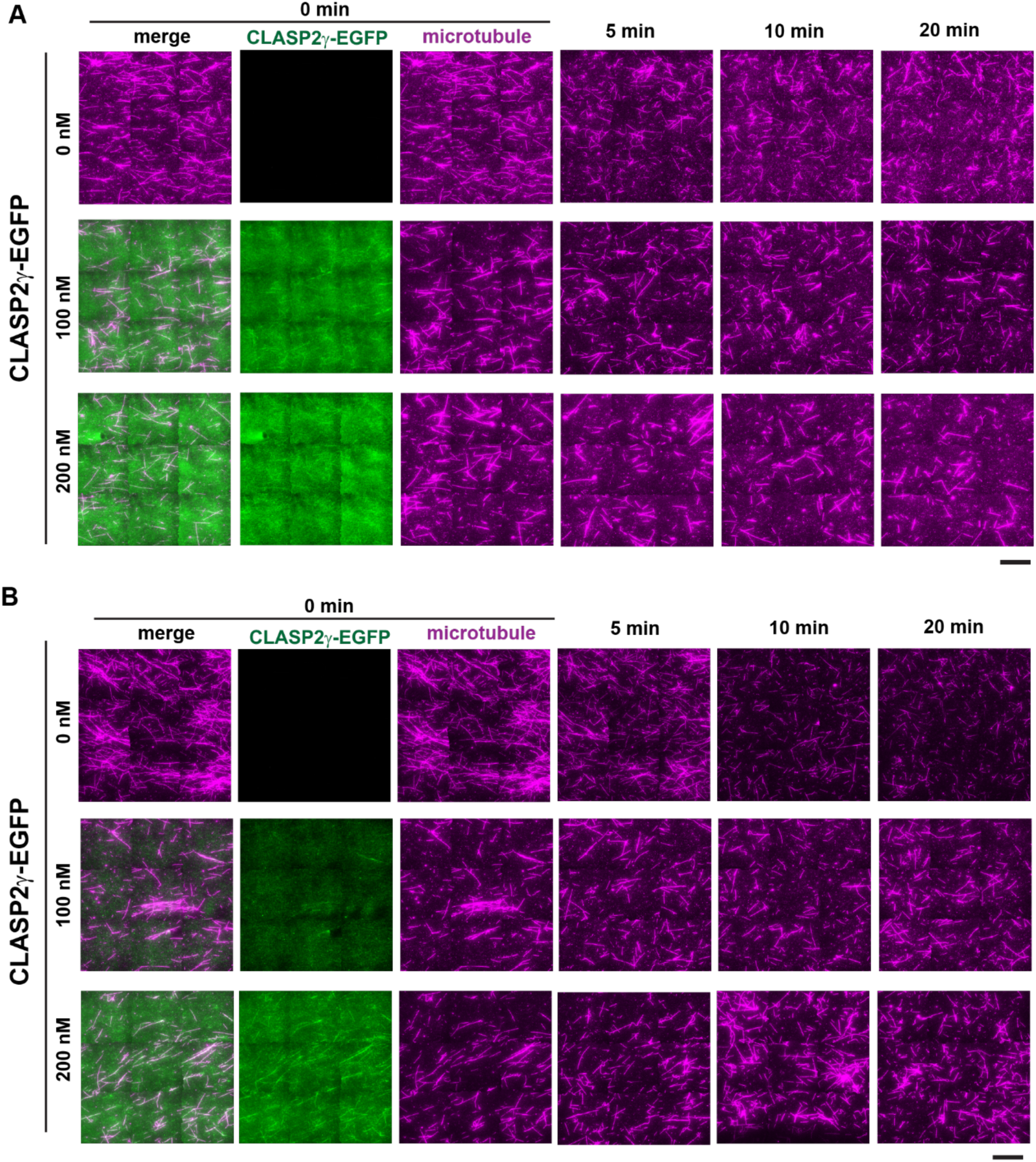
CLASP2γ protects microtubules from kinesin-induced destruction. Representative images of microtubules remaining after damage induced by (A) KIF5C(WT) or (B) KIF5C(Δ6) in the absence or presence of the indicated concentrations of CLASP2γ-EGFP. Scale bar: 20 µm.

**Fig S5.**
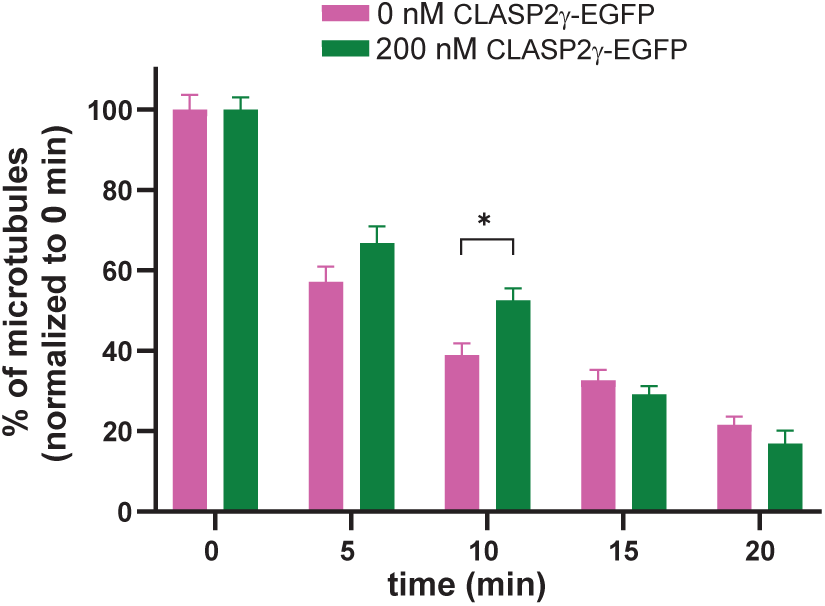
CLASP2γ does not protect microtubules from kinesin-induced damage in absence of free tubulin. Quantification of the microtubules remaining after KIF5C(Δ6)-induced microtubule damage over time in absence (magenta) or presence of 200 nM CLASP2γ-EGFP (green). The total number of microtubules was counted per field of view at 0 min (immediately after ATP addition to start KIF5C motility) and then after 5, 10, 15, or 20 min. The percent of microtubules remaining in each FOV was determined, averaged for 27 FOV across three independent experiments, and normalized to the 0 min time point (error bars: SE). * p<0.05 (two-tailed t test).

## Movie legends.

**Movie S1: KIF5C(WT)-induced microtubule destruction.** Representative movie of microtubule destruction assayswith glycerol-stabilized GDP-microtubules driven by ∼200 nM surface-bound KIF5C(WT) motors in the presence of 1 mM GTP, 2 mM ATP, and 7 µM free tubulin. Images were acquired for 5 min at 5 sec intervals.

**Movie S2: KIF5C(Δ6)-induced microtubule destruction.** Representative movie of microtubule destruction assay with glycerol-stabilized GDP-microtubules driven by ∼200 nM KIF5C(Δ6) motors in presence of 1 mM GTP, 2 mM ATP, and 7 µM free tubulin. Images were acquired for 5 min at 5 sec intervals.

## References

1. Gudimchuk, N.B. and J.R. McIntosh, Regulation of microtubule dynamics, mechanics and function through the growing tip. Nature reviews. Molecular cell biology, 2021. 22(12): p. 777–795.

2. Lawrence, E.J., S. Chatterjee, and M. Zanic, More is different: Reconstituting complexity in microtubule regulation. The Journal of biological chemistry, 2023. 299(12): p. 105398.

3. Cleary, J.M. and W.O. Hancock, Molecular mechanisms underlying microtubule growth dynamics. Current biology, 2021. 31(10): p. R560–R573.

4. Brouhard, G.J. and L.M. Rice, Microtubule dynamics: an interplay of biochemistry and mechanics. Nature reviews. Molecular cell biology, 2018. 19(7): p. 451–463.

5. Mitchison, T. and M. Kirschner, Dynamic instability of microtubule growth. Nature (London), 1984. 312(5991): p. 237-242.

6. Chew, Y.-M. and R.A. Cross, Structural switching of tubulin in the microtubule lattice. Biochemical Society transactions, 2025. 53(1): p. 161–171.

7. Mehidi, A. and C. Aumeier, Regulation of the microtubule network; the shaft matters. Current opinion in systems biology, 2023. 34: p. 100457.

8. Verhey, K.J. and R. Ohi, Causes, costs and consequences of kinesin motors communicating through the microtubule lattice. Journal of cell science, 2023. 136(5).

9. Romeiro Motta, M., S. Biswas, and L. Schaedel, Beyond uniformity: Exploring the heterogeneous and dynamic nature of the microtubule lattice. European journal of cell biology, 2023. 102(4): p. 151370.

10. Verhey, K.J., et al., The Kinesin Superfamily. 2015, Springer Netherlands: The Netherlands. p. 1–26.

11. Verhey, K.J., N. Kaul, and V. Soppina, Kinesin Assembly and Movement in Cells. Annual review of biophysics, 2011. 40(1): p. 267–288.

12. Yildiz, A., Mechanism and regulation of kinesin motors. Nature reviews. Molecular cell biology, 2025. 26(2): p. 86–103.

13. Verhey, K.J. and J.W. Hammond, Traffic control: regulation of kinesin motors. Nature reviews. Molecular cell biology, 2009. 10(11): p. 765–777.

14. Hirokawa, N., et al., Kinesin superfamily motor proteins and intracellular transport. Nature reviews. Molecular cell biology, 2009. 10(10): p. 682–696.

15. Triclin, S., et al., Self-repair protects microtubules from their destruction by molecular motors. Nature materials, 2021. 20(6): p. 883–891.

16. Andreu-Carbó, M., et al., Motor usage imprints microtubule stability along the shaft. Developmental cell, 2022. 57(1): p. 5–18.e8.

17. Budaitis, B.G., et al., A kinesin-1 variant reveals motor-induced microtubule damage in cells. Current biology, 2022. 32(11): p. 2416–2429.e6.

18. Nandakumar, S., et al., Kinesin-Induced Buckling Reveals the Limits of Microtubule Self-Repair. Advanced science, 2026. 13(26): p. e21721-n/a.

19. Bieling, P., et al., Reconstitution of a microtubule plus-end tracking system in vitro. Nature, 2007. 450(7172): p. 1100–1105.

20. Komarova, Y., et al., Mammalian end binding proteins control persistent microtubule growth. The Journal of cell biology, 2009. 184(5): p. 691–706.

21. Vitre, B., et al., EB1 regulates microtubule dynamics and tubulin sheet closure in vitro. Nature cell biology, 2008. 10(4): p. 415–421.

22. Nakamura, M., X.Z. Zhou, and K.P. Lu, Critical role for the EB1 and APC interaction in the regulation of microtubule polymerization. Current biology, 2001. 11(13): p. 1062–1067.

23. Vemu, A., et al., Severing enzymes amplify microtubule arrays through lattice GTP-tubulin incorporation. Science (American Association for the Advancement of Science), 2018. 361(6404).

24. Aumeier, C., et al., Self-repair promotes microtubule rescue. Nature cell biology, 2016. 18(10): p. 1054–1064.

25. Szczesna, J.O.S., Sarbanes, A., and Roll-Mecak, A. Tubulin flux at spastin-induced nanodamage sites regulates microtubule rescue frequency and EB1 lifetimes. Proc. Natl. Acad. Sci. U.S.A., 2026. 123(22): p. e2517683123.

26. Miesch, J., et al., Phase separation of +TIP networks regulates microtubule dynamics. Proceedings of the National Academy of Sciences - PNAS, 2023. 120(35): p. 1–12.

27. Komarova, Y.A., et al., Cytoplasmic linker proteins promote microtubule rescue in vivo. The Journal of cell biology, 2002. 159(4): p. 589–599.

28. Manna, T., et al., Suppression of Microtubule Dynamic Instability by the +TIP Protein EB1 and Its Modulation by the CAP-Gly Domain of p150Glued. Biochemistry (Easton), 2008. 47(2): p. 779–786.

29. Mishima, M., et al., Structural basis for tubulin recognition by cytoplasmic linker protein 170 and its autoinhibition. Proceedings of the National Academy of Sciences - PNAS, 2007. 104(25): p. 10346–10351.

30. Arnal, I., et al., CLIP-170/Tubulin-Curved Oligomers Coassemble at Microtubule Ends and Promote Rescues. Current biology, 2004. 14(23): p. 2086–2095.

31. Henrie, H., et al., Stress-induced phosphorylation of CLIP-170 by JNK promotes microtubule rescue. The Journal of cell biology, 2020. 219(7).

32. de Forges, H., et al., Localized Mechanical Stress Promotes Microtubule Rescue. Current biology, 2016. 26(24): p. 3399–3406.

33. Lawrence, E.J. and M. Zanic, Rescuing microtubules from the brink of catastrophe: CLASPs lead the way. Current opinion in cell biology, 2019. 56: p. 94–101.

34. Lawrence, E.J., et al., Human CLASP2 specifically regulates microtubule catastrophe and rescue. Molecular biology of the cell, 2018. 29(10): p. 1168–1177.

35. Lawrence, E.J., M. Zanic, and L.M. Rice, CLASPs at a glance. Journal of cell science, 2020. 133(8).

36. Akhmanova, A., et al., CLASPs Are CLIP-115 and -170 Associating Proteins Involved in the Regional Regulation of Microtubule Dynamics in Motile Fibroblasts. Cell, 2001. 104(6): p. 923–935.

37. Mimori-Kiyosue, Y., et al., CLASP1 and CLASP2 bind to EB1 and regulate microtubule plus-end dynamics at the cell cortex. The Journal of cell biology, 2005. 168(1): p. 141–153.

38. Aher, A., et al., CLASP Suppresses Microtubule Catastrophes through a Single TOG Domain. Developmental cell, 2018. 46(1): p. 40–58.e8.

39. Aher, A., et al., CLASP Mediates Microtubule Repair by Restricting Lattice Damage and Regulating Tubulin Incorporation. Current biology, 2020. 30(11): p. 2175–2183.e6.

40. Al-Bassam, J., et al., CLASP Promotes Microtubule Rescue by Recruiting Tubulin Dimers to the Microtubule. Developmental cell, 2010. 19(2): p. 245–258.

41. Farmer, V.J. and M. Zanic, TOG-domain proteins. Current biology, 2021. 31(10): p. R499–R501.

42. Luo, W., et al., CLASP2 recognizes tubulins exposed at the microtubule plus-end in a nucleotide state-sensitive manner. Science advances, 2023. 9(1): p. eabq5404.

43. Majumdar, S., et al., An isolated CLASP TOG domain suppresses microtubule catastrophe and promotes rescue. Molecular biology of the cell, 2018. 29(11): p. 1359–1375.

44. Biswas, S., et al., Tau accelerates tubulin exchange in the microtubule lattice. Nature physics, 2025. 21(10): p. 1616–1628.

45. Reid, T.A., et al., Structural state recognition facilitates tip tracking of EB1 at growing microtubule ends. eLife, 2019. 8.

46. Lawrence, E.J., S. Chatterjee, and M. Zanic, CLASPs stabilize the pre-catastrophe intermediate state between microtubule growth and shrinkage. The Journal of cell biology, 2023. 222(7).

47. Schmidt, T.G.M., et al., Molecular Interaction Between the Strep-tag Affinity Peptide and its Cognate Target, Streptavidin. Journal of molecular biology, 1996. 255(5): p. 753–766.

48. Schmidt, T.G.M., et al., Development of the Twin-Strep-tag® and its application for purification of recombinant proteins from cell culture supernatants. Protein expression and purification, 2013. 92(1): p. 54–61.

49. Li, Y., et al., Compressive forces stabilize microtubules in living cells. Nature materials, 2023. 22: p. 913–924.

50. Ju, R.J., et al., Compression-dependent microtubule reinforcement enables cells to navigate confined environments. Nature cell biology, 2024. 26(9): p. 1520–1534.

51. Schneider, C.A., W.S. Rasband, and K.W. Eliceiri, NIH Image to ImageJ: 25 years of image analysis. Nature methods, 2012. 9(7): p. 671–675.

52. Yue, Y., et al., Altered chemomechanical coupling causes impaired motility of the kinesin-4 motors KIF27 and KIF7. The Journal of cell biology, 2018. 217(4): p. 1319–1334.

53. Wasilko, D.J., et al., The titerless infected-cells preservation and scale-up (TIPS) method for large-scale production of NO-sensitive human soluble guanylate cyclase (sGC) from insect cells infected with recombinant baculovirus. Protein expression and purification, 2009. 65(2): p. 122–132.

